# Erythroid precursors regulate local oxygen tension and repair outcomes in the bone marrow niche

**DOI:** 10.1101/2025.01.10.632440

**Authors:** Annemarie Lang, Joseph M. Collins, Madhura P. Nijsure, Simin Belali, Mohd Parvez Khan, Yasaman Moharrer, Ernestina Schipani, Yvette Y. Yien, Yi Fan, Michael Gelinsky, Sergei A. Vinogradov, Cameron Koch, Joel D. Boerckel

**Author notes:** Correspondence: Annemarie Lang – Joel D. Boerckel –.

## Abstract

Oxygen tension dynamically regulates stem cell fate and tissue regeneration, yet how local oxygen availability is controlled within the bone marrow niche remains poorly understood. While bone marrow injury, such as by bone fracture, disrupts marrow vasculature, the consequences on local oxygen tension remain unclear. Here, we show in mice that while the tissue oxygen tension in bone marrow is low (25 mmHg, ∼4% O_2_), intracellular oxygenation is heterogeneous and erythroid cells are high in oxygen. Bone fracture elevates oxygen tension in the injured bone marrow (>55 mmHg, ∼8%), which persists for over a week post-injury. This oxygen elevation results not from angiogenesis, but rather from localized expansion of erythroid precursor cells in the injured bone marrow. The activated erythroid precursors synthesize hemoglobin and accumulate oxygen, acting as local modulators of oxygen tension. Blocking transferrin receptor 1 (CD71)–mediated iron uptake impairs hemoglobin synthesis, reduces local oxygen levels, and enhances bone regeneration through increased angiogenesis and osteogenesis. These findings identify erythroid precursors as active regulators of local oxygen availability in the bone marrow niche, which may be targetable to enhance tissue regeneration.

## Main

Tissue oxygenation varies throughout the body. Atmospheric oxygen tension, or partial pressure of oxygen (pO_2_), is about 155 mmHg, which corresponds to 21% fraction of oxygen in air. Arterial blood reaches oxygen fractions of about 14%, and organs like the spleen maintain levels around 8% ^1^. In contrast, bone marrow is among the most hypoxic tissues in the body, typically ranging between 1-4% oxygen, depending on location ^2^. This hypoxia is not pathological but rather a defining feature of the bone marrow microenvironment. Spatial oxygen gradients within the marrow give rise to distinct functional niches. The peri-sinusoidal regions, which are deeply embedded and densely vascularized, exhibit the lowest oxygen tensions, approximately 9.9 mmHg (1.3%) ^2^, while the endosteal zones, perfused by small arterioles are comparatively less hypoxic, with oxygen levels around 13.5 mmHg (1.8%) ^2^. These spatial heterogeneities regulate, for example, hematopoietic stem cell behavior. Low oxygen tensions direct metabolic reprogramming, quiescence, and self-renewal to maintain stem cell integrity and function^3^. While bone marrow oxygenation has been well characterized at homeostasis, the extent to which individual cell populations experience and regulate oxygen tension, particularly in response to bone marrow injury, remains poorly understood.

Bone marrow is composed of diverse cell types, including hematopoietic progenitors, stromal cells, and erythroid precursors, each with distinct metabolic demands and potential contributions to local oxygen dynamics. Transient perturbations, such as injury to the bone marrow niche by bone fracture, enable interrogation of how this cellular heterogeneity influences oxygen distribution within the niche during injury and regeneration. Bone fracture ruptures marrow vasculature, leading to hematoma formation and bleeding arrest. However, the impact of fracture-induced bone marrow injury on local oxygenation remains unclear^4-7^ and the contribution of marrow-resident cells to oxygen levels is unknown.

Here we use a pipeline of complementary tools to assess oxygen microenvironment dynamics in long bone marrow under homeostasis and injury contexts. We demonstrate that, in homeostasis, a majority of marrow-resident cells are hypoxic, while a minority of marrow cells, including erythroid lineage cells, exhibit higher oxygen levels. Marrow injury by bone fracture, elevates oxygen in the local microenvironment. This enrichment is driven by transient activation of local erythropoiesis, characterized by erythroid precursor expansion and hemoglobin synthesis. Mechanistically, we show that inhibition of transferrin receptor (Tfrc)-mediated iron uptake impairs heme synthesis, reduces oxygen availability, and re-establishes a hypoxic niche.

### Homeostatic bone marrow exhibits intracellular oxygen heterogeneity

To examine cellular-level oxygen heterogeneity in the bone marrow niche, we combined two orthogonal approaches: direct tissue oxygen tension measurements using phosphorescence quenching of Oxyphor PtG4 ^8-10^, and intracellular hypoxia labeling using EF5 ^11-13^. In steady-state femoral bone marrow, Oxyphor PtG4 detected oxygen tension of 28.8 ± 7.0 mmHg (∼4%; **Fig. 1a**), consistent with previous reports ^2^ and significantly lower than oxygen levels in the oxygen-rich spleen^14,15^. The bone marrow is a densely vascularized tissue composed of distinct hematopoietic and stromal compartments (**Fig. 1b**). Among them, erythroid lineage cells actively synthesize hemoglobin during development and may serve as local oxygen buffers due to their heme content. To determine whether intracellular oxygenation varies by cell type, we administered EF5, which covalently labels hypoxic cells in an inverse pO₂-dependent manner. In intact bone marrow, more than 50% of cells were EF5-positive, confirming widespread cellular hypoxia under basal conditions (**Fig. 1b, c**). In contrast, the spleen showed minimal EF5 labelling, consistent with its higher oxygen tension (**Fig. 1c**). We next compared oxygenation between erythroid and non-erythroid marrow cells (**Fig. 1d**). Flow cytometric analysis of CD71⁺Ter119⁺ erythroid precursors revealed that most were EF5-negative, while the majority of non-erythroid cells were EF5-positive, indicating higher intracellular oxygen levels in erythroid cells (**Fig. 1e; Extended Data Fig. 1**). To validate this, we cultured freshly isolated bone marrow cells under defined oxygen conditions for 1 hour and found that even under severe hypoxia (0.1% O₂), a subset of cells remained EF5-negative, suggesting intrinsic oxygen-binding capacity (**Fig. 1f; Extended Data Fig. 2**) or heterogeneity in their ability to metabolically reduce EF5. While it remains unclear whether erythroid precursors in bone marrow actively modulate niche oxygenation, their hemoglobin content may enable oxygen storage, as observed in mature erythrocytes^16,17^ and other non-erythroid, hemoglobin-expressing cells ^18 19 20 21^.

**Figure 1:**
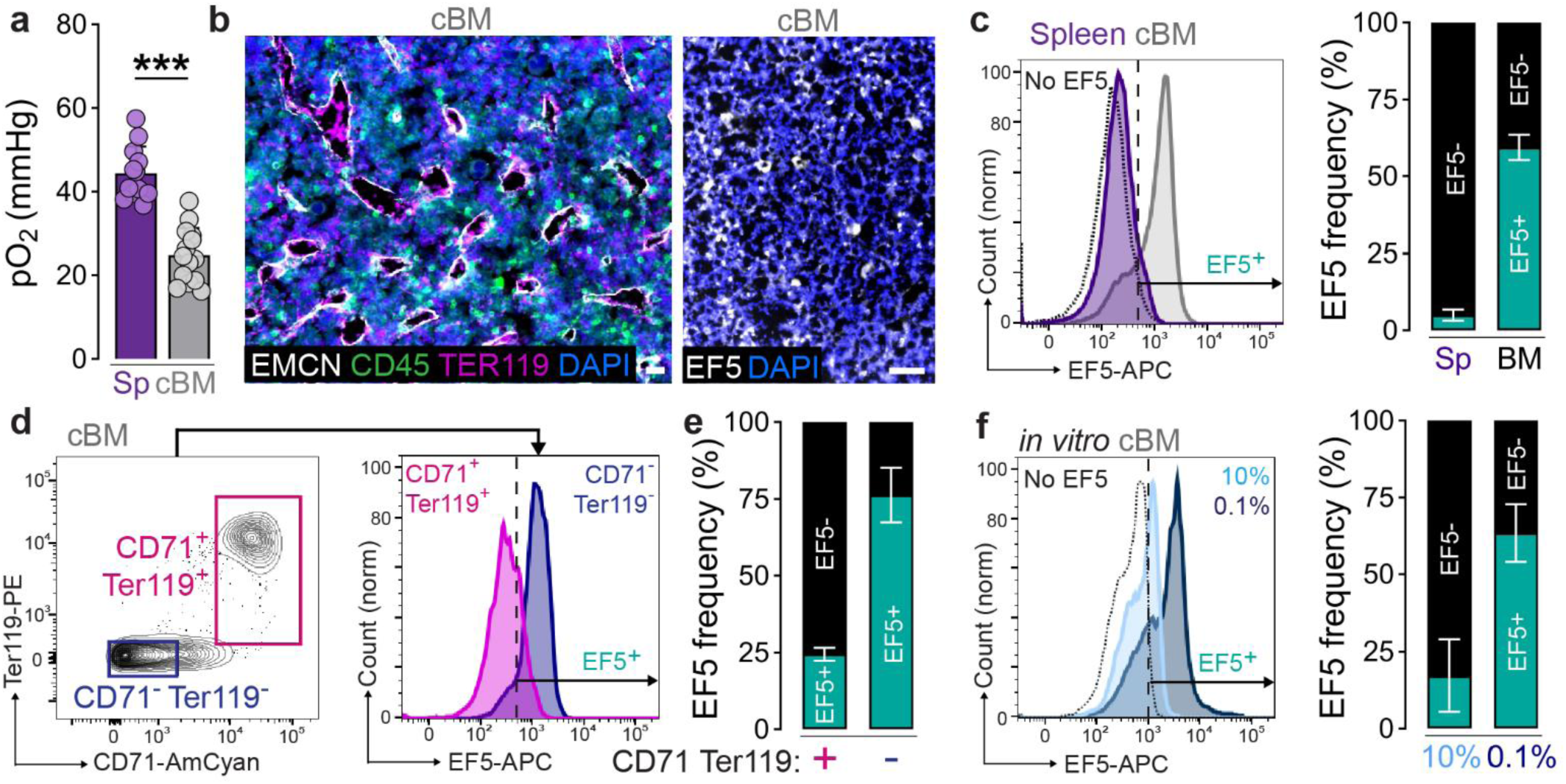
Erythroid cells in bone marrow exhibit high intracellular oxygen and may serve as local oxygen buffers. (**a**) Direct oxygen measurement in the spleen (Sp) compared to uninjured bone marrow (cBM). Data show individual data points for n = 6-14 mice at least 3 independent experiments and are mean ± s.d. *P* values were calculated using two-tailed Student’s *t*-test. \*\*\**P* < 0.001. (**b**) Immunofluorescent localization of EMCN^+^ vessel, CD45^+^ hematopoietic stem and progenitor cells (HSPCs), and TER119^+^ erythroid cells in bone marrow (left) and next to EF5 immunofluorescence staining in bone marrow (right). Scale bars, 20 µm and 50 µm, respectively. (**c**) Representative histogram and quantification of EF5 staining in bone marrow and spleen. Data are mean ± s.e.m. for n = 5-10 mice from at least 5 independent experiments. (**d, e**) Flow cytometry analysis of erythroid lineage cells (CD71^+^ Ter119^+^) compared to non-erythroid (CD71^-^ Ter119^-^) cells. Data are mean ± s.e.m. for n = 4 mice from 4 independent experiments. (**f**) Representative histogram and quantification of EF5 staining in cells isolated from the bone marrow, then cultivated for 1h at different levels of pO_2_ (represented by fractional oxygen). Data are mean ± s.e.m. for n = 4 mice from 2 independent experiments.

### Oxygen elevation at the injury site is independent of angiogenesis

To investigate how dynamic perturbations influence cellular oxygenation within the bone marrow niche, we used bone fracture as a physiological model of localized disruption. Fracture acutely injures the bone marrow and its vasculature, creating a spatially and temporally defined zone of altered perfusion. This enables in situ interrogation of how oxygen levels are regulated across distinct cell types and regions during injury and early regeneration. We hypothesized that fracture-induced vascular disruption would cause profound local hypoxia, particularly in marrow regions distal to the revascularization front (**Fig. 2a, b**). After fracture, the revascularization front proceeds from proximal to distal at the injury site, with most significant vascular network disruption distal to the fracture gap ^22,23^ (**Fig. 2c**). We reasoned that bone marrow cells distal to the fracture would be particularly and severely hypoxic after injury. To our surprise, the entire injury site at 3 days post-injury (dpi) was EF5-negative (**Fig. 2d**), indicating an absence of hypoxic cells. We next isolated cells from the injury site for quantitative flow cytometry. Bone fracture significantly reduced the frequency of hypoxic EF5-positive cells, compared to contralateral bone marrow, at 3, 5, and 7 dpi (**Fig. 2e; Extended Data Fig. 3**). We next directly measure tissue oxygen tension in uninjured bone marrow and the injury site. The injury site measured 57.9 ± 11.5 mmHg pO_2_ (∼8%) at 3 dpi, significantly higher than in the contralateral bone marrow and comparable to the spleen (**Extended Data Fig. 4a**). Oxygen levels declined over time to 30 mmHg pO_2_ (4%) by 14 dpi, comparable to contralateral bone marrow (**Fig. 2f**), consistent with EF5 staining (**Fig. 2d**). We next asked whether this unexpected elevation in oxygen could be explained by local angiogenesis. Surprisingly, oxygen tension and vascular density were inversely correlated across samples (R = -0.98; **Fig. 2g, h; Extended Data Fig. 4b-d**). Further, while blood vessel rupture releases oxygen-carrying erythrocytes into the injury site, hematoma formation rapidly stops this bleeding. Despite this, injury site oxygen tension remained elevated beyond 7 dpi. To test whether impaired angiogenesis would induce hypoxia, we applied ambulatory mechanical loading to disrupt vessel formation ^5,22^. Compliant fixation indeed disrupted vascularity, but did not induce hypoxia at the injury site (**Fig. 2i, j**). These findings reveal that the early injury site is unexpectedly oxygen-rich and that this elevation is not dependent on revascularization, suggesting an alternative, non-vascular source of local oxygen binding.

**Figure 2:**
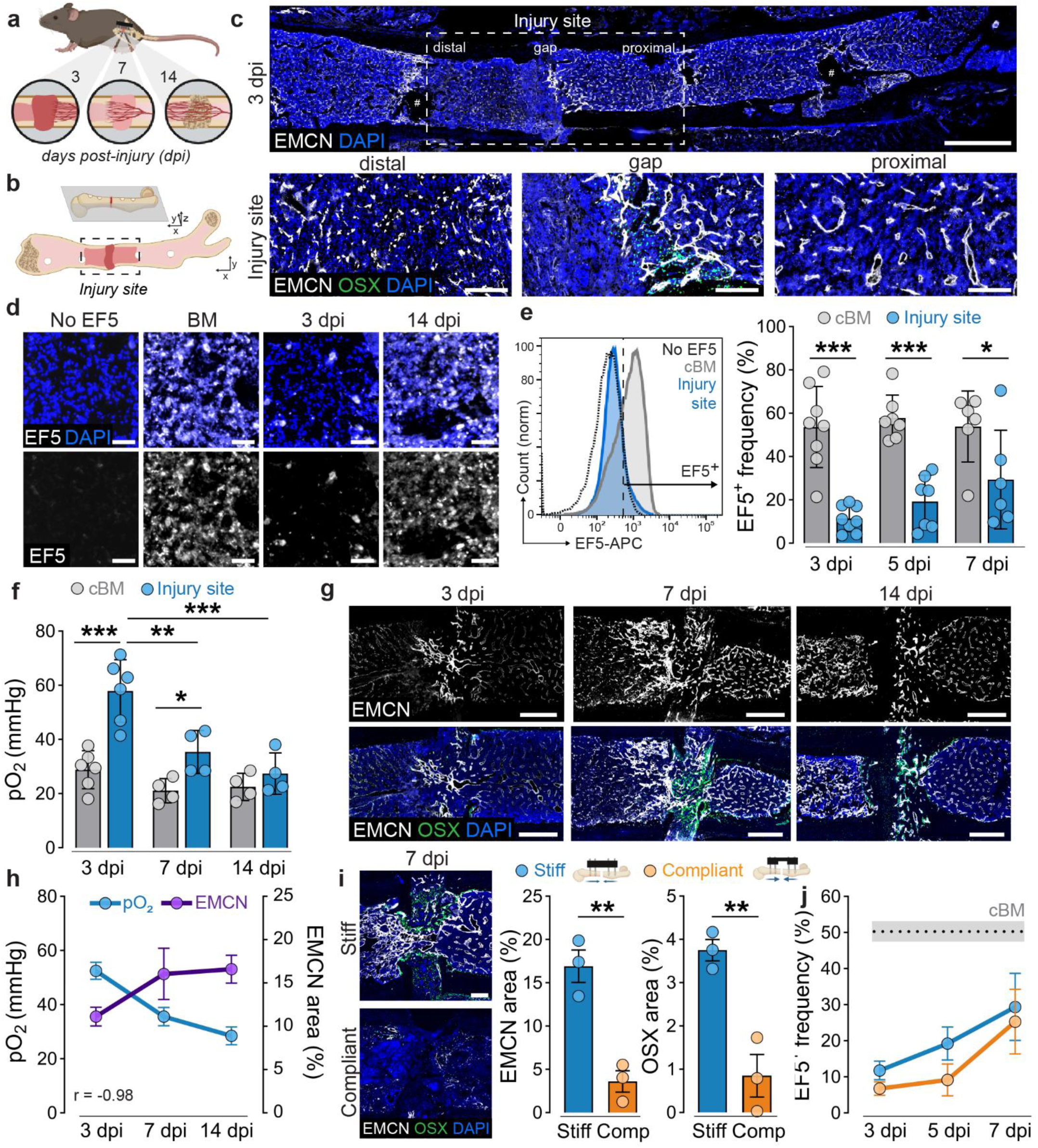
The early injury site is not hypoxic. (**a**) Schematic of the tissue regeneration process in mice over 14 days post-injury (dpi), illustrating hematoma at 3 dpi, revascularization at 7 dpi and cartilage/bone formation at 14 dpi ^24^. (**b**) We refer to the injury site as the fracture gap and adjacent bone marrow ^24^. (**c**) Immunofluorescent localization of EMCN^+^ vessel formation and OSX^+^ osteoprogenitor infiltration along the revascularization front at 3 dpi. Scale bars, 1 mm (top) and 200 µm (bottom). ^#^The center of pin holes caused by external fixator device are exemplary marked. (**d**) We injected mice with EF5 for immunolocalization of hypoxic cells in uninjured bone marrow (cBM), and the injury site at 3 and 14 dpi. Scale bars, 50 µm. Representative images from n = 3-4 mice per group in 3 independent experiments. (**e**) Flow cytometry analysis showing representative histograms and quantification of EF5^+^ cell frequency in no-EF5 control, cBM and injury site cells at 3, 5 and 7 dpi. Data show individual data points for n = 6-8 mice per group and timepoint from up to 5 independent experiments and are mean ± s.d. *P* values were calculated using two-way ANOVA with Šidák multiple comparison test at each timepoint. *P* (cBM vs. Fx) < 0.001. (**f**) We used Oxyphor PtG4 to directly measure oxygen tension (pO_2_) in cBM and at the injury site at 3, 7, and 14 dpi. Data show individual data points for n = 4-6 mice per group and timepoint from at least 2 independent experiments and are mean ± s.d. *P* values were calculated using two-way ANOVA with Tukey multiple comparison test for preselected comparisons. *P* (cBM vs. Fx; time; interaction) < 0.01. (**g**) Immunofluorescent localization of EMCN and OSX, indicating revascularization from proximal to distal. Scale bars, 200 µm. Representative images from n = 3-6 mice per timepoint in 3 independent experiments. (**h**) Quantification and correlation analysis of pO_2_ (mean of injury site) and EMCN^+^ area (total injury site) in the same samples. Data are mean ± s.e.m. for n = 4-6 mice per group and timepoint from at least 2 independent experiments. Pearson correlation coefficient: R = -0.98, *P* = 0.12. (**i**) Immunofluorescent localization of EMCN and OSX at 7 dpi under stiff and compliant fixation ^24^. Scale bars, 400 µm. Data show individual data points for n = 3 mice per group and are mean ± s.e.m. *P* values were calculated using two-tailed Student’s *t*-test. \**P* < 0.05, \*\**P* < 0.01 and \*\*\**P* < 0.001. (**j**) Quantification of EF5^+^ cell frequency under stiff and compliant fixation. Data are mean ± s.e.m for n = 4-7 mice per group per timepoint from 5 independent experiments.

### Bone marrow erythroid precursor cells expand after injury

Bone marrow contains resident hematopoetic stem and progenitor cells (HSPCs) that create new red blood cells through erythropoiesis. We used single cell RNA-sequencing to identify cell populations in contralateral bone marrow and the injury site at 3 dpi (**Fig. 3a**). We identified eight major cell clusters by canonical marker expression (**Fig. 3b, c; Extended Data Fig. 5**). Innate immune cells, including monocytes, macrophages and granulocytes, represented the majority of cells at the injury site, consistent with known inflammation kinetics ^23,25,26^ (**Extended Data Fig. 5**). Other cell types included HSPCs, adaptive immune cells (T- and B-cells) and erythroid cells (**Extended Data Fig. 6**). Relative to contralateral bone marrow, bone injury upregulated genes involved in oxidative phosphorylation across all cell types, particularly in HSPCs, neutrophils, and erythroid cells (**Extended Data Fig. 7**). Erythroid-lineage cells bind oxygen by iron-containing hemoglobin (**Fig. 3d**) and are characterized by expression of the transcription factor *Gata1*, the membrane protein *Glyphorin A (Gypa)*, and the iron importer *Transferrin receptor (Tfrc)* ^27-29^ (**Fig. 3c**). Hemoglobin synthesis in erythroid precursors includes production of heme. Heme is synthesized through multiple enzymatic steps in the cytosol and mitochondria, culminating in the insertion of Fe^2+^ ion into the porphyrin ring^30^ (**Fig. 3d**). Simultaneously, globin proteins are produced in the cytoplasm (**Fig. 3d**). We found bone marrow erythroid precursor cells at the injury site, marked by expression of genes responsible for the heme synthesis and adult globin (**Fig. 3e,f**). Gene set enrichment analysis suggests that bone injury induced erythroid precursor proliferation at the injury site (**Fig. 3g**). To test whether injury activates bone marrow erythroid cells, we performed secondary Louvain clustering on the erythroid cell cluster. We identified seven erythroid cell states, which we assigned to three different developmental stages based on lineage-specific gene expression profiles ^31^: megakaryocytic-erythroid progenitors (MEP), proerythroblasts (ProE), and erythroblasts (EryB) (**Fig. 3h, i**). Cells from the injury site contained a significantly higher frequency of ProE cells compared to contralateral bone marrow, as measured by both scRNA-seq (**Fig. 3j**). Thus, bone marrow injury may amplify erythroid precursor cells at the injury site, which could influence local oxygen concentration.

**Figure 3:**
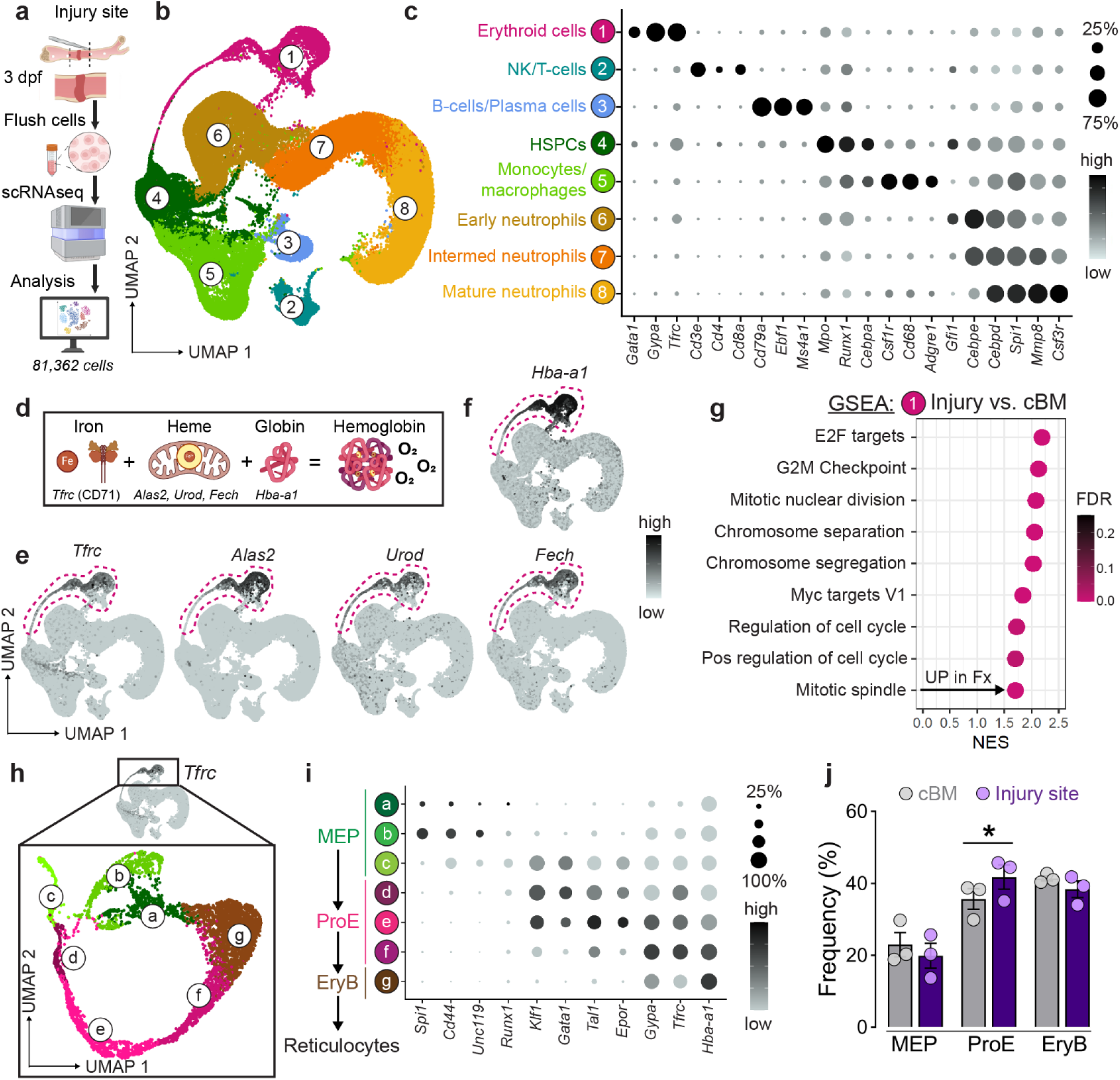
Erythroid cells express molecular machinery for oxygen binding. (**a**) Schematic illustrating scRNA-sequencing of cells flushed from the injury site at 3 dpi (Fx) or the contralateral bone marrow (cBM). We removed dead cells and lysed mature red blood cells, but did not digest the samples or sort prior to scRNA-seq ^24^. (**b**) UMAP and Louvain clustering. Integrated data sets from n = 3 mice per group from 3 independent experiments. (**c**) Cell type annotations based on canonical marker expression. (**d**) Schematic illustration of hemoglobin synthesis ^24^ and (**e, f**) feature plots of relevant genes on the UMAP. (**g**) Gene set enrichment analysis (GSEA) highlighting upregulated pathways and processes in erythroid cells at the injury site compared to cBM. (**h**) Secondary Louvain clustering of erythroid cells. Integrated data sets from n = 3 mice per group from 3 independent experiments. (**i**) Erythroid cell differentiation state annotations based on developmental stage-specific marker expression. (**j**) Frequency of cells in the different clusters normalized to the absolute cell count per cluster. Data show individual data points from n = 3 mice per group from 3 independent experiments and are mean ± s.e.m. *P* values were calculated using two-tailed paired Student’s *t*-test.

### Bone marrow injury induces local erythropoiesis

To determine whether erythroid expansion at the injury site reflects a true erythropoietic response, we quantified erythroid precursor populations by flow cytometry ^32^ (**Fig. 4a,b; Extended Data Fig. 8a**). Injury significantly increased the frequency of early-stage erythroid cells, including ProE, in the injured bone marrow. In parallel, we observed a reduction in late-stage EryB and an increase in peripheral blood reticulocytes at 3 dpi, (**Fig. 4c; Extended Data Fig. 8a**). To test whether this response was systemically regulated, we measured circulating levels of erythropoietin (Epo), the key hormone driving erythropoiesis. Serum Epo levels remained unchanged at 3 dpi (**Fig. 4d**), suggesting that the erythropoietic response was locally restricted rather than systemically induced. To spatially visualize this local erythropoiesis, we immunostained for CD71 and Ter119-expressing erythroid cells. Injury increased erythroid cell abundance adjacent to the injury site, but the contralateral (i.e., opposite limb) and ipsilateral bone marrow (i.e., same limb but far from the injury site) exhibited similar, homeostatic levels of CD71^+^ Ter119^+^ erythroid cells (**Fig. 4e**). Injury site-adjacent CD71^+^ erythroid cells declined in abundance from 3 to 14 dpi (**Fig. 4f; Extended Data Fig. 8b-d**), paralleling the decrease in O_2_ levels (cf. **Fig. 2h**). These findings indicate that bone marrow injury induces a transient and spatially localized erythropoietic response. This local activation, independent of systemic Epo signaling, may contribute to the transient elevation in oxygen tension observed during early tissue regeneration.

**Figure 4:**
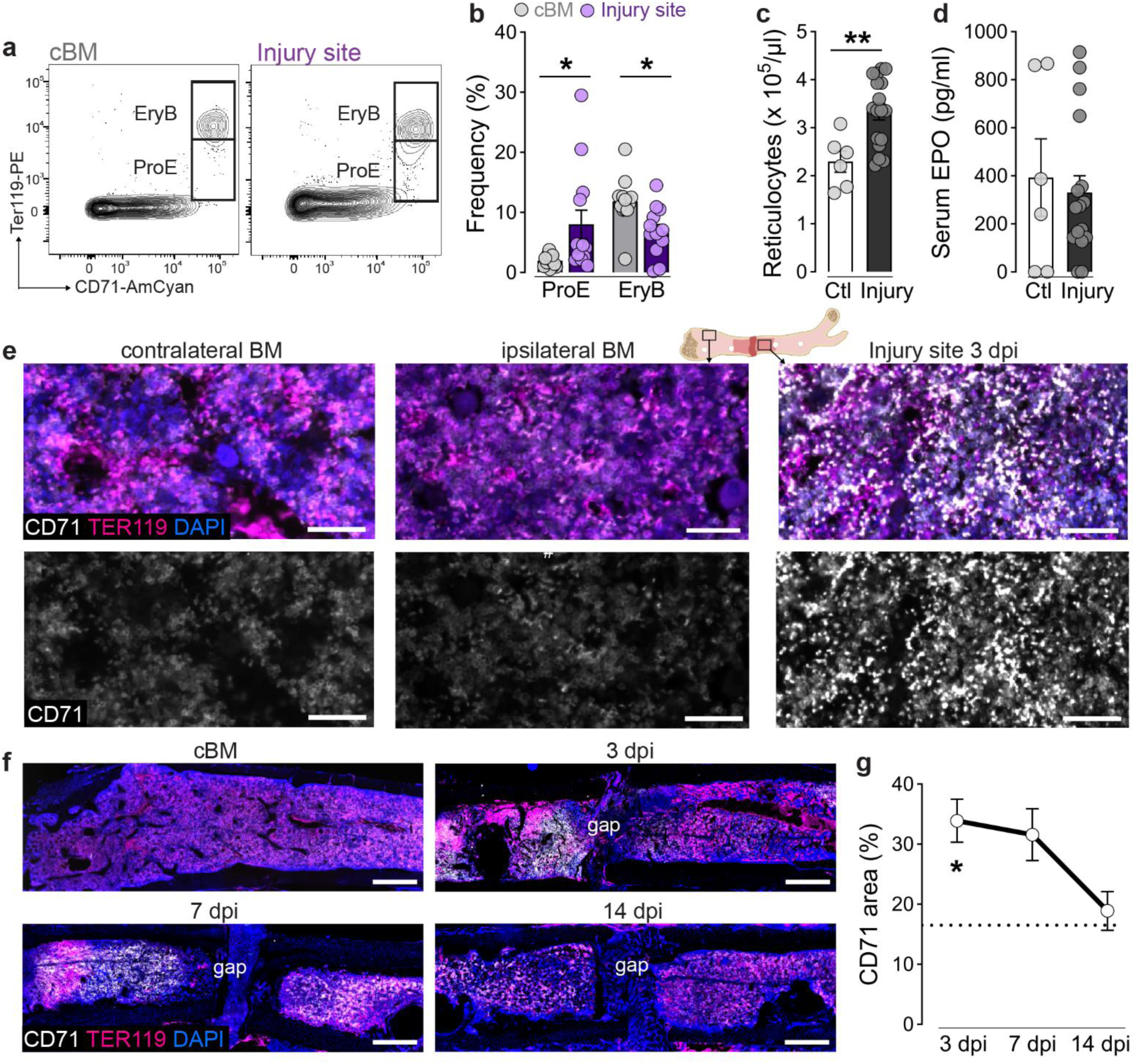
Bone marrow injury activates local erythropoiesis. (**a**) Flow cytometry analysis and representative histograms of ProE and EryB based on Ter119 marker intensity. (**b**) Frequencies of ProE (Ter119^med^ CD71^high^) and EryB (Ter119^high^ CD71^high^). Data show individual data points from n = 13 mice from more than 5 independent experiments and are mean ± s.e.m. *P* values were calculated using one-way ANOVA with Šidák multiple comparison test for preselected pairs. (**c**) Concentration of reticulocytes in peripheral blood. Data show individual data points from n = 6-17 mice from more than 5 independent experiments and are mean ± s.e.m. *P* values were calculated using two-tailed Student’s *t*-test. (**d**) Serum concentration of Erythropoietin (EPO) in peripheral blood. Concentration of reticulocytes in peripheral blood. Data show individual data points from n = 6-17 mice from more than 5 independent experiments and are mean ± s.e.m. *P* values were calculated using two-tailed Student’s *t*-test. (**e**) Immunolocalization of Ter119 and CD71 positive cells in contralateral and ipsilateral bone marrow and at the injury site ^24^. We refer to the ipsilateral bone marrow as the bone marrow regions in the injured bone but distant from the injury site. Scale bars, 50 µm. Representative images for n = 4-8 mice per timepoint from 3 independent experiments. (**f**) Representative images of Ter119 and CD71 positive cells in the cBM and at the injury site at 3, 7 and 14 dpi. Scale bars, 500 µm. (**g**) Quantification of CD71 positive area as sum of distal and proximal injury site over time. Dashed line indicates normal bone marrow control. Data are mean ± s.e.m. for n = 4-8 mice per timepoint from 3 independent experiments. *P* values were calculated using one-way ANOVA with Tukey multiple comparison test. \**P* < 0.05, and \*\**P* < 0.01.

### Erythroid precursors concentrate oxygen at the injury site

Next, we sought to determine how local erythropoiesis increased oxygen at the injury site and the impact on tissue regeneration. During their maturation, erythroid precursor cells import large quantities of iron to support hemoglobin formation and oxygen binding ^33,34^. The primary iron importer in erythroid cells is Transferrin receptor 1 (CD71, encoded by *Trfc*) (**Fig. 3c, e; Extended Data Fig. 9a**). Therefore, to block iron import in cells at the injury site, we locally injected a CD71-blocking antibody (or isotype control IgG) immediately after injury (**Fig. 5a**). CD71 blockade had no effect on the increase in erythroid cell numbers at the injury site (**Fig. 5b; Extended Data Fig. 9b**), but, in erythroid cells, significantly downregulated the expression of genes required for heme synthesis (**Fig. 5c; Extended Data Fig. 9c**). Further, CD71 blockade significantly reduced oxygen at the injury site, measured by EF5 and independently by Oxyphor PtG4 (**Fig. 5d, e; Extended Data Fig. 9d**), returning oxygen tension in the injury site at 3 dpi to the level of native bone marrow (∼5%; **Fig. 5f**). Importantly, in erythroid cells at the injury site, CD71 blockade significantly increased EF5-positivity, establishing that CD71 blockade decreased oxygenation of the erythroid precursors per se (**Fig. 5f; Extended Data Fig. 9e,f**). Further, CD71 blockade significantly enhanced bone formation (**Fig. 5h, i; Extended Data Fig. 10**), osteoprogenitor activation, and angiogenesis at the injury site at 14 dpi (**Fig. 5j, k**). Together, these findings suggest that injury-activated erythroid precursors concentrate oxygen at the injury site, but preventing erythroid cell hemoglobin production induces local hypoxia and enhances tissue regeneration.

**Figure 5:**
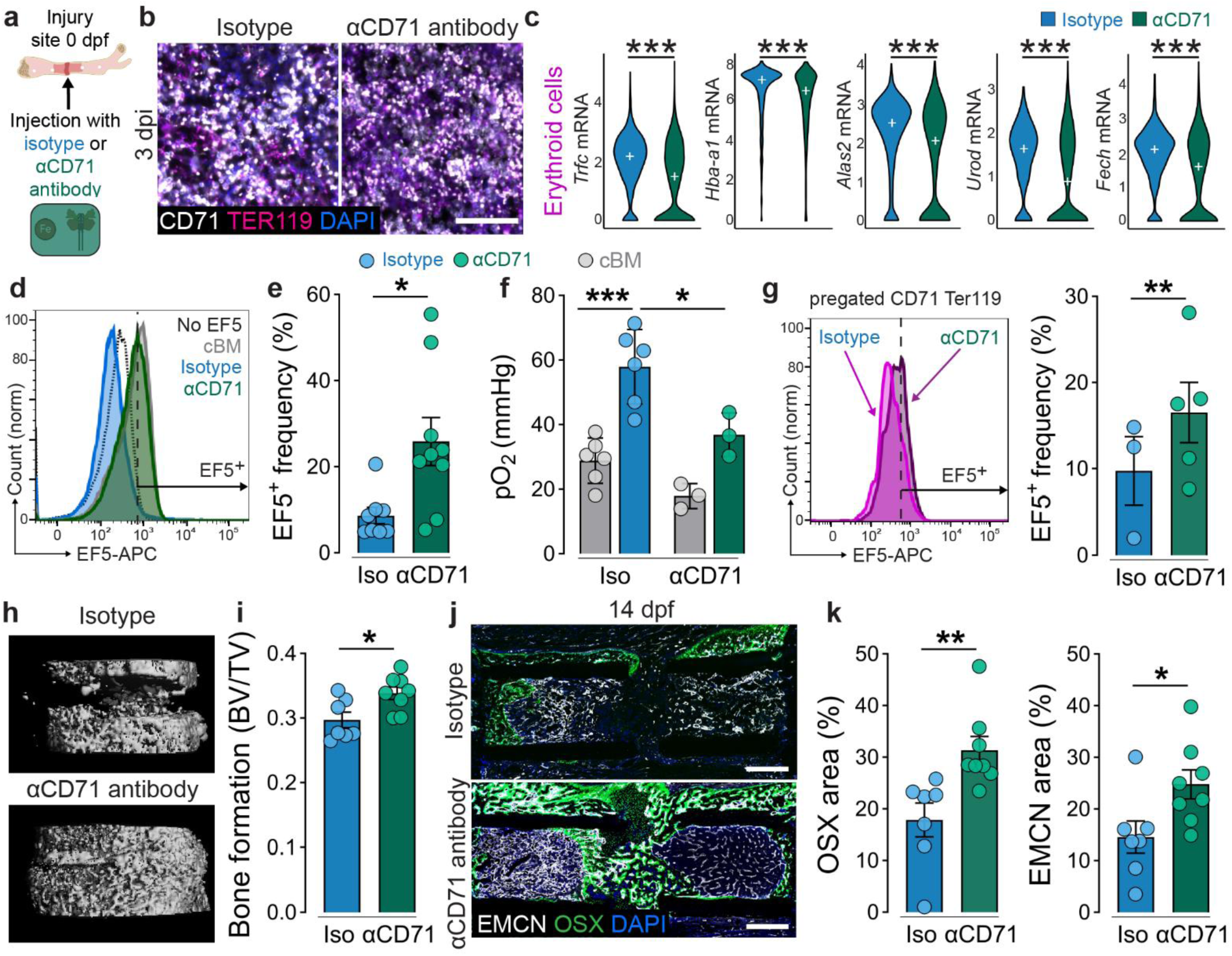
CD71 blockade induces bone marrow niche hypoxia and promotes tissue regeneration. (**a**) CD71 blocking antibody, or isotype control IgG, were injected into the injury site immediately after injury ^24^. (**b**) Immunolocalization of Ter119 and CD71 positive cells at the injury site at 3 dpi after αCD71 antibody treatment. Scale bar, 50 µm. Representative image for n = 3 mice from 3 independent experiments. (**c**) Erythroid cell expression of genes involved in heme synthesis. Integrated data sets from n = 3 mice per group from 3 independent experiments. White cross indicates median. *P* values were calculated using MAST with Bonferroni corrections. *P_adj_*: *Trfc* = 2.3E-46; *Hba-a1* = 9.8E-29; *Alas2* = 3.0E-41; *Urod* = 8.0E-46; *Fech* = 1.0E-40. (**d**) Flow cytometry analysis of EF5 frequency displaying representative histogram and (**e**) quantification of EF5^+^ cell frequency in cells isolated from the injury site at 3 dpi. Data show individual data points for n = 8-9 mice per group from 5 independent experiments and are mean ± s.e.m. *P* values were calculated using two-tailed Student’s *t*-test. (**f**) Direct measurement of oxygen tension in the fracture gap and cBM at 3 dpi. Data for Iso are reproduced from Fig. 1f. αCD71 data show individual data points for n = 3 mice from 3 independent experiments and are mean ± s.d. *P* values were calculated using one-way ANOVA with Šidák multiple comparison test for preselected comparisons. (**g**) Flow cytometry analysis of EF5 positivity in erythroid lineage cells (CD71^+^ Ter119^+^) at 3 dpi. Data show individual data points for n = 3-5 mice per group from 3 independent experiments and are mean ± s.e.m. *P* values were calculated using two-tailed paired Student’s *t*-test. (**h, i**) MicroCT analysis of bone formation (bone volume/total volume) at 14 dpi. (**j, k**) Immunofluorescence analysis of EMCN^+^ vessel formation and OSX^+^ progenitor infiltration at the injury site at 14 dpi. Scale bars, 500 µm. Data show individual data points for n = 9-10 mice per group from 2 independent experiments and are mean ± s.e.m. *P* values were calculated using two-tailed Student’s *t*-test. \**P* < 0.05, \*\**P* < 0.01 and \*\*\**P* < 0.001.

## Conclusions

Here, we identify a previously unrecognized mechanism of oxygen regulation within the bone marrow niche. Using bone fracture as a physiological model of bone marrow injury, we show that the early injury site features elevated oxygen, driven by local activation of erythroid precursor cells. These cells accumulate in the injured bone marrow, upregulate hemoglobin synthesis, and modulate oxygen within the niche. Blocking iron import via transferrin receptor impairs hemoglobin production, reduces oxygen availability, and enhances tissue regeneration. This work positions erythroid precursors as a targetable cellular component within the bone marrow microenvironment and establishes fracture as a model to study oxygen-regulatory mechanisms in regenerative contexts (**Extended Data Fig. 11**).

The effects of bone fracture on bone marrow oxygen tension are poorly understood.^35-39^. Here, we show using multiple orthogonal methods that bone marrow injury by bone fracture induces elevated oxygen in the injured bone marrow. Oxygen tension at the fracture site was first reported by Brighton and Krebs in 1972 ^36^. Using the rabbit fibula as a model, they measured the fracture hematoma to be hypoxic (6.3 mmHg). However, the rabbit fibula contains minimal bone marrow and minimal erythropoietic activity ^40^. Most long-bone injuries involve the bone marrow, and our data are consistent with recent findings from a sheep tibial osteotomy model ^37^.

Mature red blood cells carry oxygen throughout the body, but whether immature erythroid progenitor/precursor cells are also capable of binding oxygen is poorly understood. Here, we show that bone marrow-resident erythroid precursor cells can influence their local oxygen environment and that this is functionally relevant in the context of tissue regeneration. We do not yet know the molecular mechanisms by which bone injury activates local erythropoiesis. Injury activation of erythropoiesis occurs locally, without systemic change in EPO, but coincides with macrophage mobilization to the injury site. In the bone marrow, erythropoiesis is regulated under homeostasis and stress by specialized macrophages in erythroblastic islands ^41,42^. Therefore, we speculate that localized erythropoiesis in the bone marrow could be dependent on local macrophage activation ^26,43^.

Blood vessels and angiogenesis are critical to tissue regeneration. Disruption of blood vessel formation, whether genetic, pharmacologic, or mechanical alters the course of tissue regeneration, particularly stimulating cartilaginous callus formation ^22,44-47^. We and others previously attributed this chondrogenesis to reduced oxygen transport ^22,44^. However, here we found that while compliant fixation induced cartilage callus formation ^48^, mechanical disruption of angiogenesis did not induce injury site hypoxia. This suggests that elevated local pO_2_ is not due to local angiogenesis. This is consistent with a recent study demonstrating that disruption of neovascular ingrowth during bone repair induces chondrogenesis because of limited nutrient supply, particularly of fatty acids, rather than limited oxygen ^49^. Further, our observation that injury upregulated genes involved in oxidative phosphorylation in cells of the injury site, suggests an increase in the respiratory activity and makes it more likely that the elevated pO_2_ in the injured bone marrow is caused not by a decrease in cellular oxygen consumption but by oxygen concentration by erythroid cells.

We speculate that hemoglobin-producing erythroid precursor cells function as a local oxygen sink, creating an oxygen gradient as oxygen diffuses from surrounding vascularized tissues toward the injury site. This transiently maintains high oxygen tension at the injury site as long as oxygen diffusion outpaces consumption, but blockade of erythroid iron transport prevents formation of the oxygen sink and resultant oxygen gradient. Consistently, we observed an inverse relationship between injury site oxygenation and local vascular density, characterized by decreasing oxygen with blood vessel invasion. We speculate that the recovery of bone marrow hypoxia promotes angiogenesis, allowing for maturation and extravasation of the erythroid cells, clearing the injury site.

These findings may also change how we think about the evolution of fracture repair. Producing a high oxygen environment that delays regeneration seems evolutionarily counterproductive. We speculate a trade-off between making a maximally hospitable environment for healing vs. fighting microbial infection ^50,51^, during the window of greatest infection risk after fracture. Thus, high oxygen at the fracture site may prioritize pathogen control at the expense of delayed regeneration. Together, our work identifies an unexpected role of erythroid precursors in regulating oxygenation of the bone marrow niche, which may be targetable to promote regeneration.

## Methods

### Mouse femoral fracture model

Female or male C57BL/6J mice were obtained from The Jackson Laboratory at age 10 weeks and used within 2-6 weeks. All procedures adhered to IACUC regulations (University of Pennsylvania; protocol no: 806482). Veterinary care and husbandry were provided by University Laboratory Animal Resources (ULAR) at the University of Pennsylvania, following contemporary best practices. Mice were housed in a semi-barrier facility with a 12/12-hour light/dark cycle (light from 7:00 a.m. to 7:00 p.m.), a room temperature of 72 ± 2°F, and a humidity of 50 ± 10%. Food and tap water were available *ad libitum*. Mice were randomly paired per cage, which contained wooden chips, Enviro-dri, and shredded paper towels as bedding and nesting material. Additional enrichment included a clear mouse transfer tube (Braintree Scientific), a mouse double swing (Datesand Group), and a Shepherd Shack (Shepherd Specialty Papers) with an enlarged entrance to prevent injuries ^52^. The transfer tube and double swing were removed post-surgery to reduce injury risk. Mice were handled with the transfer tube, and cages were changed weekly.

*Mouse femoral fracture model:* Mice were induced with isoflurane (2–3% in 100% oxygen) and placed on a 37 °C heating pad. Anesthesia was maintained at 1.5–2% with a nose cone. Eye ointment, physiological saline (0.9% sodium chloride; 0.5 ml, s.c.) and clindamycin (45 mg/kg, s.c.) were administered. Buprenorphine SR-Lab (1 mg/kg, s.c.; Wedgewood Pharmacy) or Ethiqa XR (extended release buprenorphine; 3.25 mg/kg, s.c.; Fidelis Animal Health) were used for analgesia. The left femur was shaved and disinfected with iodine solution and 70% ethanol. A longitudinal incision was made between the knee and hip, and the musculus vastus lateralis and musculus biceps femoris were separated to expose the femur. Two external fixators (stiff: 18.1 N/mm; compliant: 3.2 N/mm ^53^, RISystem) were used to modulate ambulatory load transfer. The fixator bar was aligned parallel to the femur, and pins were inserted after pre-drilling. A 0.5 mm fracture gap was created using a Gigli wire saw (0.44 mm; RISystem) and flushed with saline. Muscle and skin were sutured in two layers, and the wound was treated with antibiotic cream. Mice were returned to their home cages, placed under an infrared lamp, and monitored until fully recovered. Diet Gel was provided on the cage floor to ensure post-surgery food and water intake. Mice were closely monitored and scored for the first 4 days, then on days 7 and 10, before euthanasia. Scoring included a composite score based on the mouse grimace score (eyes and ears only), clinical score, and a model-specific score for limping and dragging, following established systems ^54,55^. Humane endpoints were predefined, including criteria such as wound dehiscence, poor grooming, sunken eyes, hunched posture, periprosthetic fracture, severe axial deviation, lack of food and water intake, significant weight loss, bloody feces, severe respiratory issues, debilitating diarrhea, seizures, paresis, and abscesses. No humane endpoints were reached during the study. Mice were euthanized at different timepoints post-surgery using CO_2_ and cervical dislocation. Samples were collected as described below. All analyses were performed blind to the groups (fixation, treatment), with de-blinding only after all analyses were completed to avoid bias.

### EF5 cell staining

To identify hypoxic cells, we used EF5, which, as an inverse function of oxygen, is reduced by nitroreductase enzymes to form cytoplasmic EF5-protein adducts that can be identified by immunostaining.

*In vivo experiments:* Mice were injected with the hypoxia marker EF5 (10 µl/gm of 10 mM solution) and euthanized using CO_2_ and cervical dislocation after 2h. As a negative control for in situ EF5 labelling of cells with known high pO_2_, we performed EF5 staining of the spleen (nominal pO_2_ of 8-10%). As a positive control for *in sit*u EF5 labelling of marrow cells in low oxygen conditions, mice were similarly injected with 10 mM EF5 but euthanized after 30 min. The body was then kept at 37°C for 45 min (termed euthanized animal control, EAC) to allow all EF5 to be biochemically reduced in place. To control for any nonspecific background staining, we also included bone marrow cells from non-EF5 treated animals (EF5^-^).

For subsequent EF5 staining on frozen sections, fractured femora were cryo-embedded (Tissue-Tek OCT) without previous fixative treatment and stored at -80 °C until further use. For flow cytometry, cells were flushed from the injury site (i.e., region between the inner pins), the adjacent bone marrow of the same limb and the bone marrow from the uninjured contralateral bone. Flushed cells were immediately fixed with 4% paraformaldehyde for 1h on ice.

*EF5 staining on cryosections:* Consecutive 7 μm-thick sections were prepared using cryotape (Sectionlab). Sections were fixed onto glass slides and stored at -80 °C until staining. Before staining, sections were fixed in 4% paraformaldehyde, blocked with 5% goat serum, and stained 4h with Anti-EF5 antibody, clone ELK3-51 Cyanine 3 conjugate dissolved in PBS/3% BSA/0.1% Tween 20 (stock: 2 mg/mL; 1:25 to get 80 µg/ml). Sections from positive and negative controls were used to adjust imaging settings.

*In vitro experiments:* Bone marrow cells were flushed from femora of C57BL/6J female mice and transferred into culture medium (MEM + 5% FCS supplemented with HEPES and bicarbonate as buffer system and 100 µM EF5 compound). Cells were cultured without CO_2_ in aluminum chambers (shaken at ∼ 1Hz) for 2h at 37°. The gas phase in the chambers contained 10%, 2%, 0.5%, 0.1% fraction of oxygen (pO_2_). Cells were rinsed, then fixed with 4% paraformaldehyde for 1h on ice.

*EF5 staining and flow cytometry*: Fixed cells were washed with PBS and stained over night with Anti-EF5 Antibody, clone ELK3-51 Antibody, Cyanine 5 conjugate (stock: 2 mg/mL; 1:200 to get 10 µg/ml) dissolved in PBS/1.5% BSA/1.5% non-fat dry milk/0.3% Tween 20/5% mouse serum at 4°C under constant shaking. Cells were carefully washed twice with PBS/0.3% Tween 20 and once with PBS (30 min each). In case of additional staining, antibodies were diluted in PBS/2% FCS/0.3% Tween 20 and incubated on the cells for 30 min before additional washing steps and secondary fixation (PBS/1% PFA). The following antibodies were used: CD71-APC (BioLegend; clone: RI7217; catalogue number: 113819; 1:100); Ter119-PE (BioLegend; catalogue number: 116207; 1:100); CD71-Brilliant Violet 510 (AmCyan; BioLegend; catalogue number: 113823; 1:100); CD45-Brilliant Violet 510 (AmCyan; BioLegend; catalogue number: 103137; 1:200). Non-EF5 treated and no-antibody controls were conducted for every experiment. Cells were analyzed using a FACSCanto II flow cytometer (BD Biosciences) and the FlowJo software (V10).

### Immunofluorescent staining

Femora were fixed for 6h in 4% paraformaldehyde at room temperature and were then transferred into 10% ethylenediaminetetraacetic acid (EDTA) pH 7.4 for 3 days at 4 °C followed by treatment with 30% sucrose solution for 2 days before being cryo-embedded using Tissue-Tek OCT. Consecutive sections of 7 μm were prepared using cryotape (Sectionlab). Sections were fixed onto glass slides and stored at -20 °C until staining. Cryosections were rehydrated in PBS. Blocking solution (10% goat serum/PBS) was added for 30 min and antibodies were diluted in PBS/0.1% Tween20/5% goat serum. The following primary antibodies and secondary antibodies were used (incubation 2h at room temperature): EMCN (Santa Cruz; clone V.5C7; catalogue number: sc-65495; 1:100), OSX (Abcam; catalogue number: ab209484; 1:100), CD71-APC (BioLegend; clone: RI7217; catalogue number: 113819; 1:100); Ter119-PE (BioLegend; catalogue number: 116207; 1:100); all secondary antibodies were purchased from Thermo Fisher Scientific and used at an 1:500 dilution for 2h at room temperature: goat anti-rat A488 (A-11006) and goat anti-rabbit A647 (A-27040). DAPI (1 µg/mL) was added during the last washing step and sections were covered with Fluoromount-GT. Images were taken with an AxioScan and image quantification was performed using the Fiji/ImageJ software. The area of interest was manually assigned with the built-in ROI-Manager and determined with the thresholding tool.

### *In vivo* oxygen measurement

Oxygen measurements *in vivo* were performed by the phosphorescence quenching method ^8^ and the probe Oxyphor PtG4 ^9,10^. Overall, our measurements closely resembled phosphorescence quenching oximetry experiments performed previously in a variety of tissue types, including recent measurements in the brain ^56^, intestine ^57^ and tumours ^58^. Oxyphor PtG4 (50 µl, 50 µM; if necessary, dilution was done with saline) was injected in the tail vein 1 day prior to surgery to allow the probe to distribute evenly through tissues, ensuring that it would be present in the fracture hematoma and gap after osteotomy. Osteotomy surgery was performed as described above. At 3, 7 and 14 dpi, mice were anesthetized with isoflurane (2–3% in 100% oxygen) and placed on a heating pad (37°C). As anesthesia may influence tissue oxygen levels, it was maintained at the level of 2% isoflurane in 100% oxygen (flow rate 1.5 L/min), delivered using a nose cone. For analgesia, Buprenex was injected s.c. (0.1 mg/kg), and an eye ointment was applied. The fractured and contralateral bones were carefully dissected and exposed. The measurement times at the fractured and contralateral bones were kept similar. Oxygen levels were measured at different locations of the bone using a fiber-optic phosphorometer (Oxyled, Oxygen Enterprises). In brief, excitation was performed using a laser diode (630 nm), focused into a spot ∼200 μm in diameter on the bone surface. 630 nm light is able to diffuse into tissue reaching several mm’s (up to 1cm) depths. Thus, our measurements probed average pO_2_ throughout the bone under the excitation spot with dominant contributions from the regions closest to the surface. The phosphorescence was collected using a single mode plastic fiber (4 mm in diameter), which carried the light to an avalanche photodiode (Hamamatsu 12703). Excitation pulses were 10 μs-long, and the phosphorescence decays were collected and digitized (500 kHz frequency) during 500 μs after each pulse. 100 excitation/collection cycles were averaged to achieve high signal-to-noise ratios. Thus obtained phosphorescence decays were fitted to single-exponentials, and the corresponding decay times (lifetimes) were converted to pO_2_ values using a calibration obtained in independent experiments, as described previously ^10^. Animals were immediately euthanized after the measurements by cervical dislocation under deep anesthesia.

### Blood analysis and Epo ELISA

Blood was intracardially collected in deep anesthesia (CO_2_) before cervical dislocation. Samples for hematology were collected in EDTA tubes and transferred to the Clinical Pathology Laboratory, PennVet for complete blood count (CBC) analysis using an IDEXX ProCyte Dx Hematology Analyzer. Serum was collected in tubes with coagulation activator gel and stored at -20 °C. For assessing the serum Epo concentration, we used a Mouse Erythropoietin/EPO DuoSet ELISA kit (R&D Systems) following the manufacturer’s instructions.

### Antibody experiment and microCT

We injected either a monoclonal rat anti-mouse CD71 antibody (clone: 8D3; 100 µg) or a rat IgG2a isotype control (both Bio X Cell) in the fracture gap after skin closure. EF5 staining and Oxyphor measurement were performed at 3 dpi (days pos-injury) as described above. MicroCT and immunofluorescence analyses were performed at 14 dpi. Ex vivo microCT scanning was conducted using a µCT 45 desktop scanner (Scanco Medical AG) following the removal of the external fixator. Bones were secured in plastic pipettes to preserve the integrity of the callus tissue. Scanning focused on the region between the inner two pins with an isotropic voxel size of 10.4 μm (55 kVp, 72 μA, AL 0.5 mm, 1x400 ms), aligning the scan axis along the femoral diaphysis. 3D reconstruction and subsequent analyses were carried out with the accompanying software package, utilizing a global threshold of 240 mg HA/cm^3^. An individual VOI was determined for each sample, confined to the area between the middle pins. The analysis excluded the original cortical bone, focusing solely on newly formed bone.

### Single-cell RNA sequencing

*Cell isolation and preparation:* Femoral fracture surgeries were performed as described above. Cells were isolated from the following samples: intact contralateral femora and fractured femora at 3 dpi, with or without CD71 antibody treatment (n = 3 mice per group). Cells were flushed from the injury site (between middle pins) or the intact bone marrow (contralateral femur) in cold PBS and transferred through a cell strainer. Red blood cells were lysed with a Red Blood Cell Lysis kit and dead cells were then removed with a Dead Cell Removal kit (both Miltenyi Biotec) following the manufacturer’s instructions. Samples were confirmed to have > 95% viability (based on trypan blue staining). Cells were resuspended in PBS/0.4% BSA and transferred on ice to the sequencing core facility.

*Single-cell RNA sequencing and analysis.* Next-generation sequencing libraries were created utilizing the 10x Genomics Chromium Single-cell 3’ Reagent kit v3 following the manufacturer’s protocols. Each library was uniquely indexed with the Chromium dual Index Kit, pooled, and sequenced on an Illumina NovaSeq 6000 in a paired-end, dual indexing run, aiming for ∼ 21,092 reads per cell. Data processing involved the Cell Ranger pipeline (10x Genomics, v.6.1.2) for demultiplexing, aligning reads to the mm10 transcriptome, and generating feature-barcode matrices. Subsequent analysis employed Seurat v4.0 ^59^. Cells were ignored if they had fewer than 200 genes, more than 6000 genes, or over 5% mitochondrial reads. The data from individual samples underwent LogNormalization, and 2000 variable features were identified for each sample. Seurat’s alignment method was used for sample integration, followed by data scaling with ScaleData. Principal component analysis (PCA) was performed for linear dimensional reduction. The entire 81,362-cell dataset was visualized in two dimensions using UMAP, and clustering was performed with Louvain clustering. Dimensions were 1:7 and a resolution of 0.1 was applied in the clustering analysis. The FindAllMarkers function and canonical marker genes helped identify major cell types. Louvain clustering resulted in 8 clusters, which were assigned to the respective cell populations. For gene set enrichment analysis (GSEA), a pre-ranked list of genes was generated based on fold changes between comparisons and analyzed using GSEA v4.3.3. The murine hallmark gene set collection and the GO Biological Process ontology from the Molecular Signatures Database (MSigDB) was used for comparisons. Significant inter-cluster gene expression differences were identified using MAST with Bonferroni corrections. Erythroid cells constituted 1 of the initial 8 clusters. These cells were then sub-divided and reanalyzed using the pipeline described above. Iterative clustering of the erythroid cells revealed 7 clusters, which were assigned to 3 cell states based on canonical marker expression and relevant literature.

### Statistical Analysis

Power analysis was performed *a priori* for key outcome measures. Statistical analysis was conducted using GraphPad Prism version 9. To assess the Gaussian distribution, the D’Agostino-Pearson omnibus normality test was employed, along with tests for homoscedasticity. One-way ANOVA with multiple comparison tests (Tukey or Šidák) or two-tailed Student’s *t*-test were used to determine statistical significance, with a *P*-value of less than 0.05 considered significant. Sample sizes are indicated in figure legends and graphs display individual data points as scatterplots. Data are presented with error bars representing the mean ± s.e.m. or mean ± s.d. Exclusion of samples or data occurred only in verified cases of technical error.

## Acknowledgments

The authors would like to thank all members of the Boerckel lab for constructive discussions.

## Author contributions

A.L. and J.D.B. conceived and supervised the research. A.L., J.D.B., M.N., M.P., Y.M. designed and performed animal experiments. A.L. and J.M.C. performed analysis. A.L. and C.K. performed EF5 experiments and analysis. A.L., S.B. and S.A.V. performed direct oxygen measurement and analysis. M.P., E.S., Y.Y., Y.F., M.G. supported with methods, technical guidance and advice. A.L. and J.D.B. wrote the paper. All authors discussed and revised the manuscript.

## Data and materials availability

All data are available in the main text or the supplementary materials. The raw datasets used and/or analyzed during the current study are available from the corresponding author. The sequencing data discussed in this publication have been deposited in NCBI’s Gene Expression Omnibus and are accessible through GEO Series accession number GSE230260.

---

**Extended Data Figure 1:**
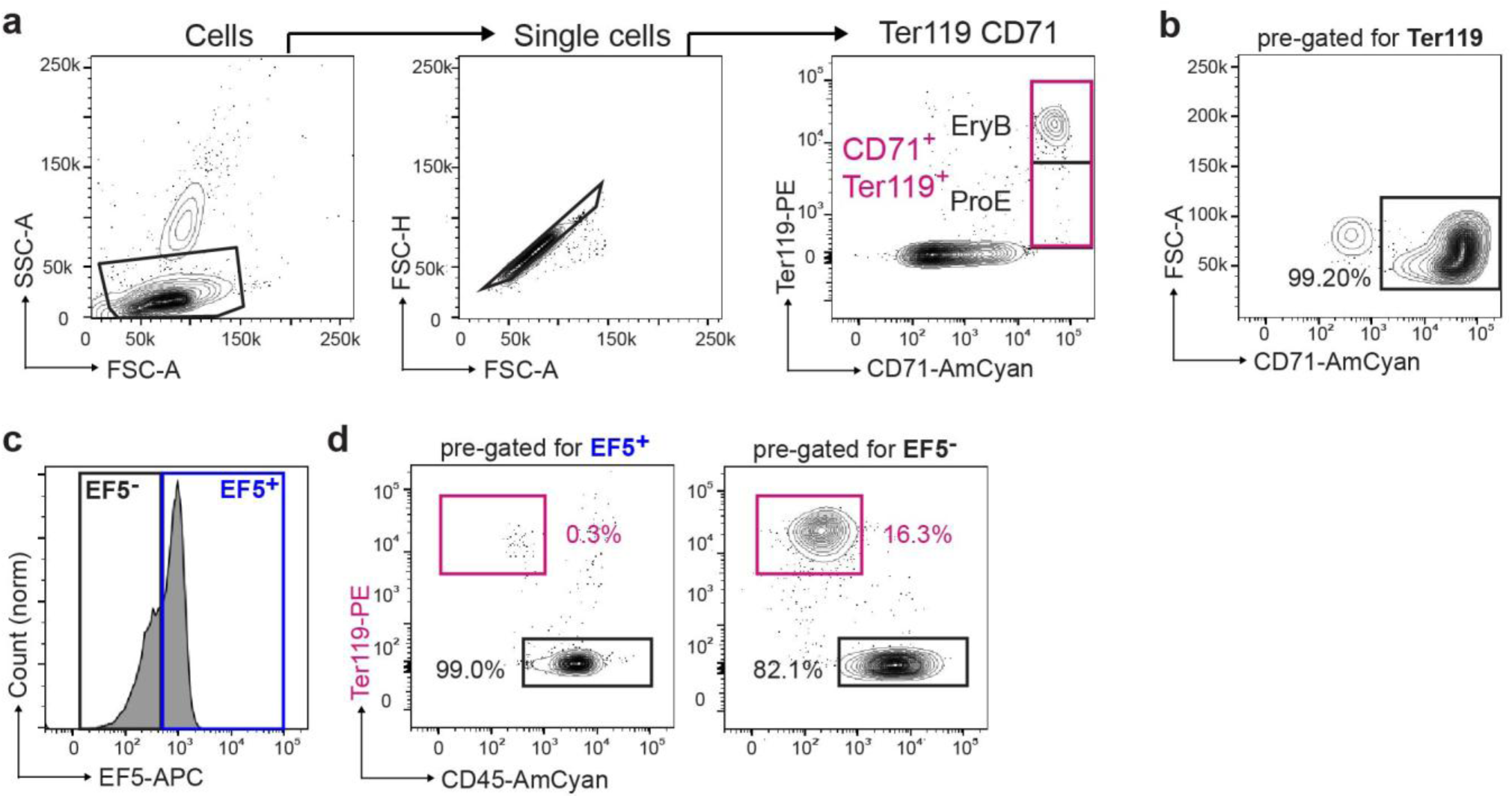
Gating and analysis strategy for CD71^+^ Ter119^+^ erythroid cells in the bone marrow. (**a**) Gating strategy for analysis of CD71^+^ Ter119^+^ cell frequency. (**b**) Detailed gating on Ter119^+^ cells confirming positive dual staining for CD71. (**c**) Uninjured bone marrow contains both EF5^+^ and EF5^-^ cells. Representative histogram of EF5 staining in bone marrow to indicate gating for EF5^-^ and EF5^+^ cells. (**d**) Of EF5^+^ cells, only 0.77 ± 0.5% were Ter119^+^ CD45^-^ erythroid cells, while of the EF5^-^ cells, 28.1 ± 8.5% of EF5^-^ cells were of erythroid lineage (mean ± s.d. for n = 4 mice from 4 independent experiments).

**Extended Data Figure 2:**
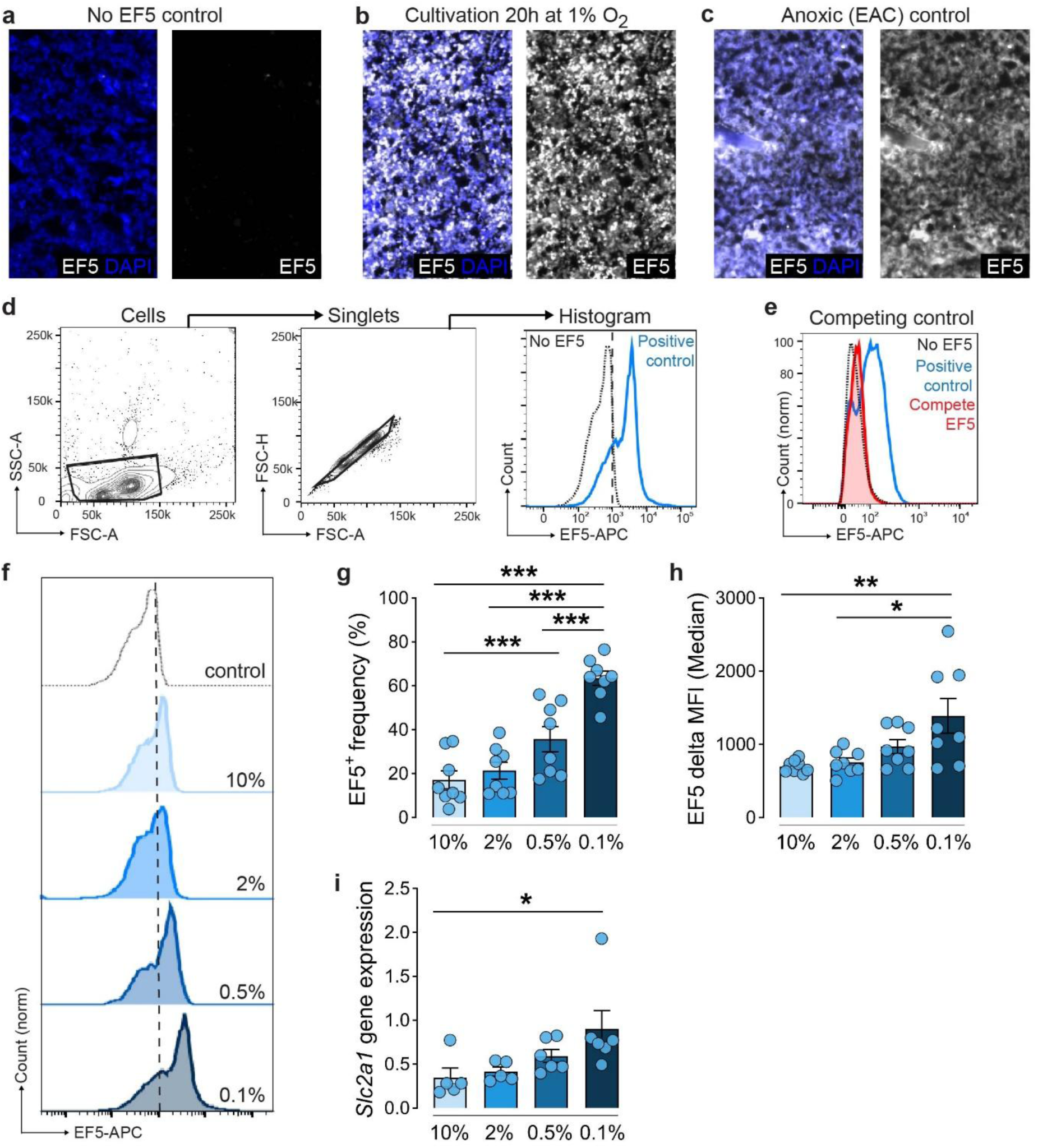
EF5 staining marks hypoxic cells. To identify hypoxic cells, we injected mice with EF5, which, as an inverse function of intracellular oxygen, is reduced by nitroreductase enzymes to form cytoplasmic EF5-protein adducts that can be identified by immunostaining. To setup the staining process for cryosections, different controls were used: (**a**) no EF5 control, (**b**) murine femora were cultivated ex vivo for 20h at 1% (fraction of oxygen) with adding EF5 compound during the last 2h and (**c**) mice were injected with EF5 compound and euthanized after 30 min. The body was then kept at 37°C for 45 min allowing tissue metabolism to create severe hypoxia and reduce all EF5 in situ (euthanized animal control, EAC). Representative images from n = 3-4 mice per group in 3 independent experiments. (**d**) Gating strategy for analysis of EF5^+^ cell frequency. (**e**) Competed-stain control. EF5 compound (which competitively inhibits Ab staining to EF5 cellular adducts) was added to the staining antibody mixture and shortly pre-incubated before being added to the isolated cells. Representative histogram for n = 2 in 2 independent experiments. (**f**) Cells isolated from the bone marrow were cultivated for 1h at different levels of pO_2_ (represented by fractions of oxygen) to confirm staining specificity as indicated by the (**g**) EF5^+^ cell frequency, (**h**) EF5 delta median fluorescence intensity and (**i**) expression of hypoxia-relevant gene. Data show individual data points for n = 4 mice from 2 independent experiments and are mean ± s.e.m. *P* values were calculated using one-way ANOVA with Šidák multiple comparison test for preselected pairs. \**P* < 0.05, \*\**P* < 0.01 and \*\*\**P* < 0.001.

**Extended Data Figure 3:**
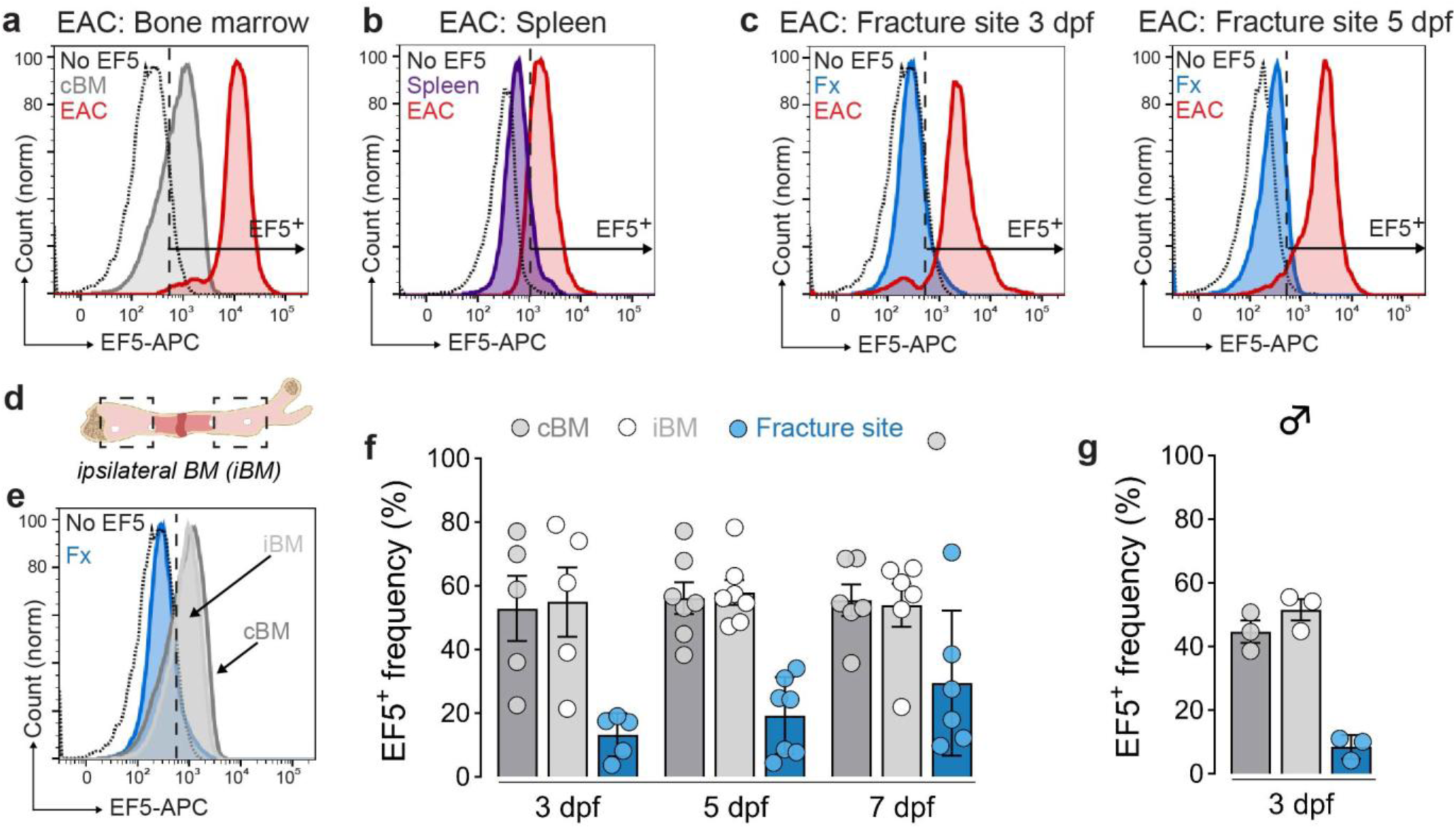
EF5 staining marks hypoxic cells at the injury site. To verify EF5 distribution and capacity to mark hypoxic cells at the injury site, we performed a control experiment to induce anoxia. Mice were injected with EF5 compound and euthanized after 30 min. The body was then kept at 37°C in anoxic conditions for 45 min (euthanized animal control, EAC). (**a-c**) EAC controls in bone marrow, spleen and the injury site 3 and 5 dpi demonstrate EF5 distribution in the tissue and EF5 dynamic range in vivo. Representative histograms from n = 2 mice in 2 independent experiments. (**d**) We refer to the ipsilateral bone marrow (iBM) as the bone marrow regions in the injured bone but distant from the injury site. (**e**) Flow cytometry analysis of EF5 staining displaying representative histogram of non-EF5 staining control, iBM, cBM, and injury site and (**f**) quantification of EF5^+^ cell frequencies at 3, 5 and 7 dpi. These experiments were performed in female mice. Data show individual data points for n = 5 mice from at least 3 independent experiments and are mean ± s.e.m. (**g**) Male mice exhibit equivalent response. Data show individual data points for n = 3 mice and are mean ± s.e.m.

**Extended Data Figure 4:**
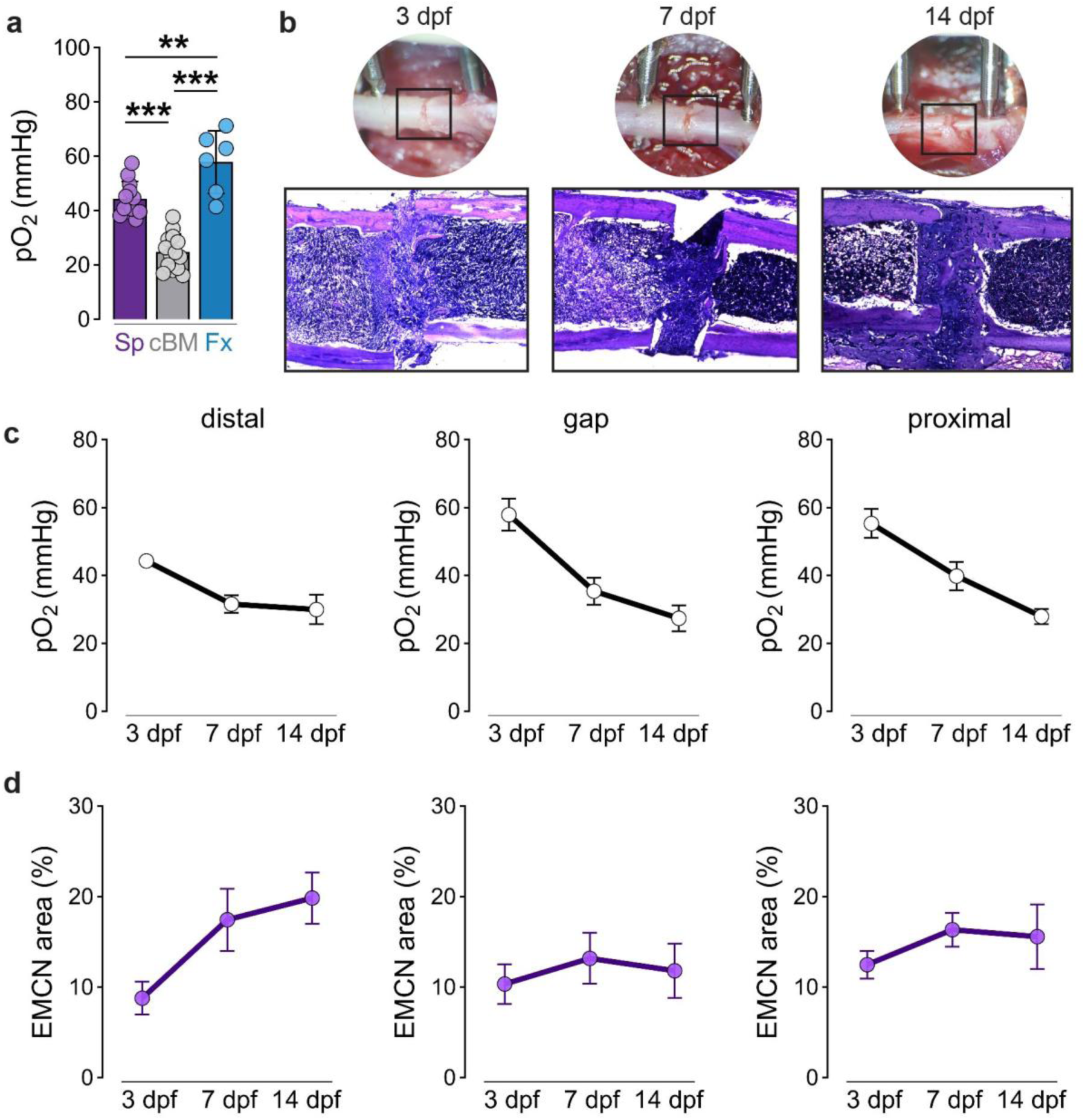
Direct oxygen measurement at the injury site. (**a**) Direct oxygen measurement in the spleen (Sp) compared to cBM and injury site at 3 dpi. Data show individual data points for n = 6-14 mice at least 3 independent experiments and are mean ± s.d. *P* values were calculated using one-way ANOVA with Tukey multiple comparison test. \*\**P* < 0.01 and \*\*\**P* < 0.001. (**b**) Macroscopic views on the injury site at 3, 7 and 14 dpi and HE staining to display cellular changes at the injury site. Representative images from for n = 4-6 mice per group and timepoint from at least 2 independent experiments. (**c**-**d**) Separate displays of temporal development of (**c**) pO_2_ values and (**d**) EMCN^+^ areas in the distal, gap and proximal injury site. Data are mean ± s.e.m. for n = 4-6 mice per group and timepoint from at least 2 independent experiments.

**Extended Data Figure 5:**
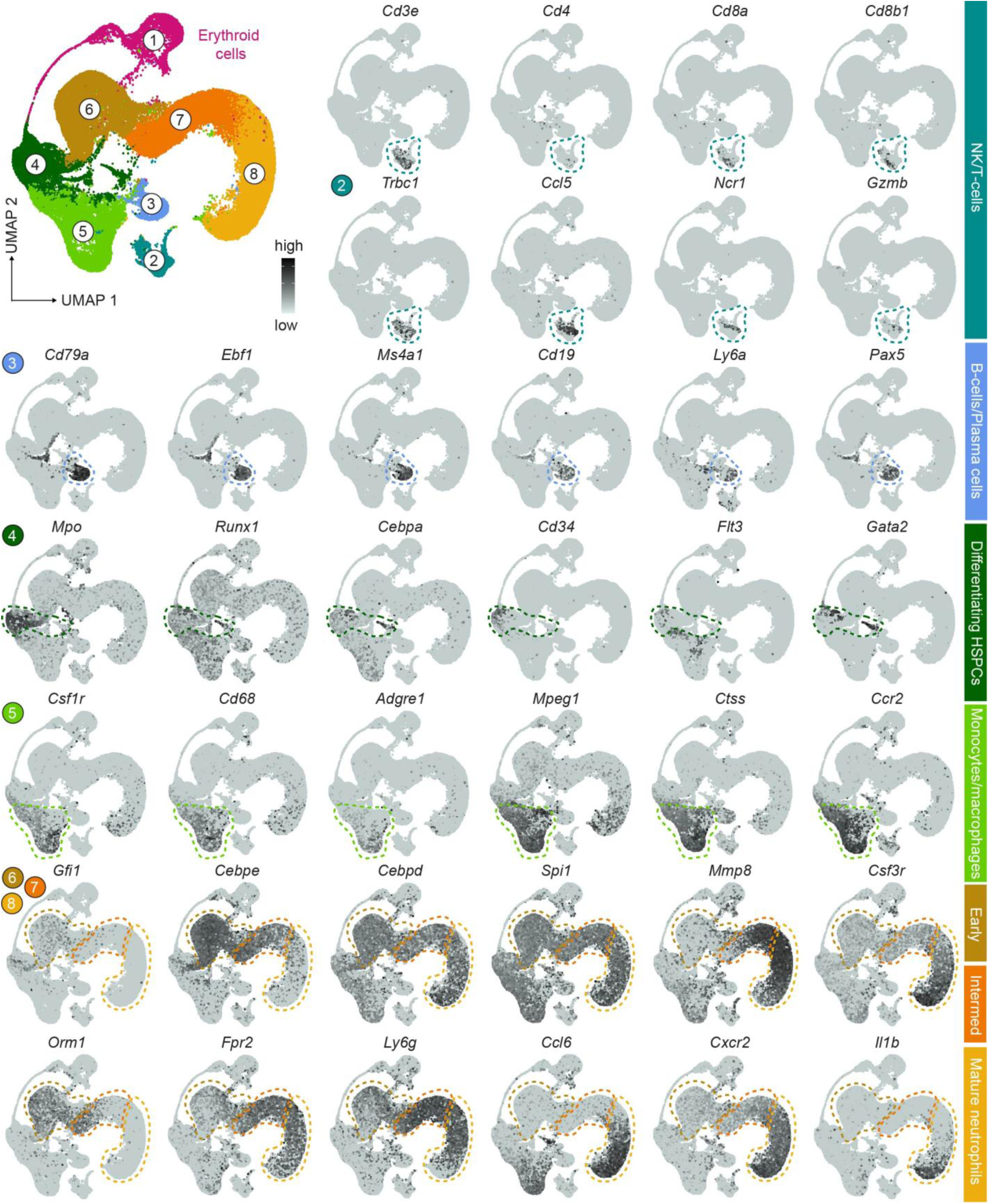
Canonical marker gene expression to identify major cell populations at the injury site at 3 dpi. Louvain clustering resulted in 8 clusters, which were assigned to the respective cell populations. Feature plots indicate relevant genes on the UMAP.

**Extended Data Figure 6:**
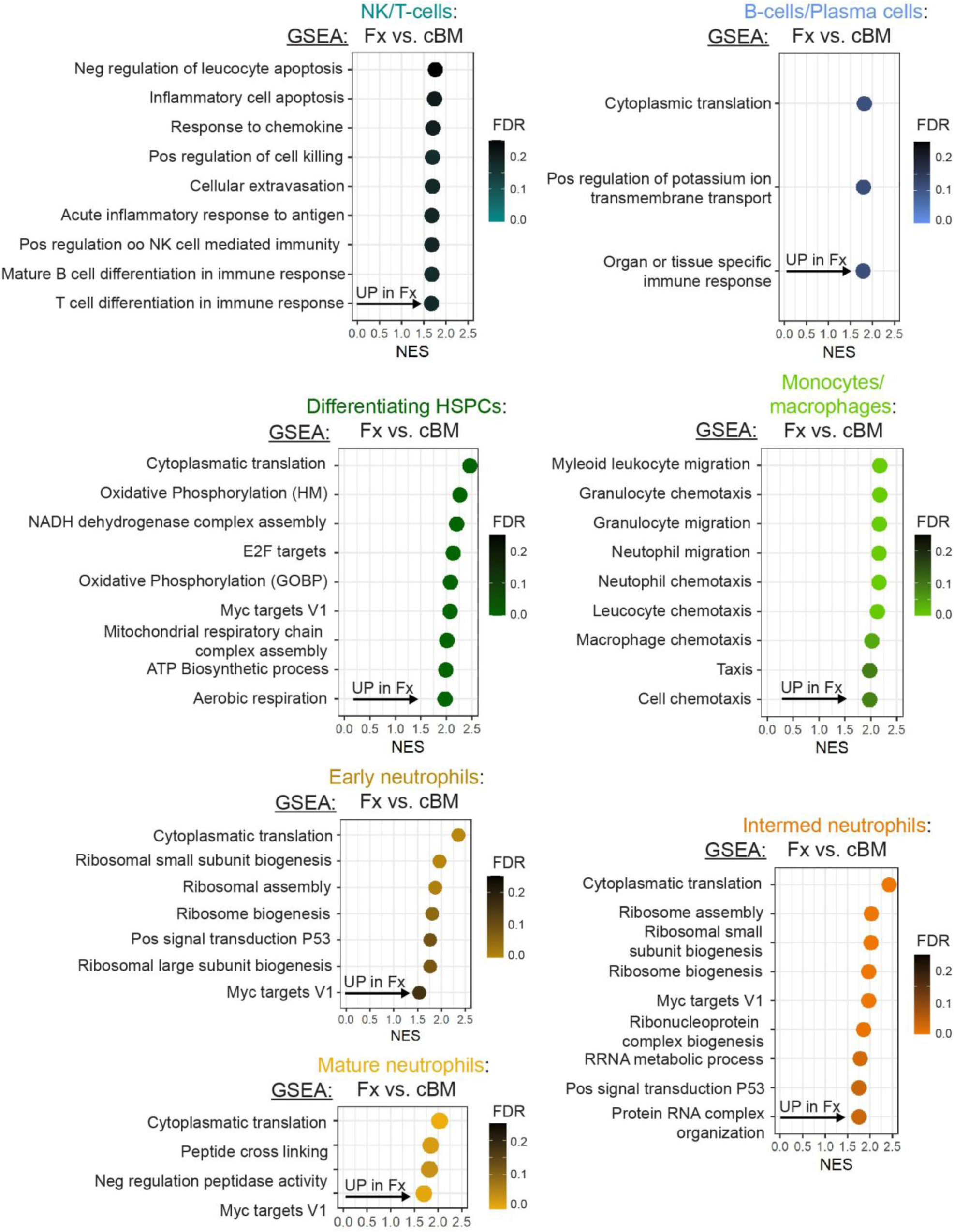
Gene set enrichment analysis (GSEA) for major cell populations at the injury site at 3 dpi. A pre-ranked list of genes was generated based on fold changes between comparisons. The murine hallmark gene set collection and the GO Biological Process ontology from the Molecular Signatures Database (MSigDB) was used for analysis. We identified activation of HSPCs and adaptive immune cells (T- and B-cells) at the injury site. Further, innate immune cells, including monocytes, macrophages and granulocytes showed pathway enrichment of known inflammation kinetics.

**Extended Data Figure 7:**
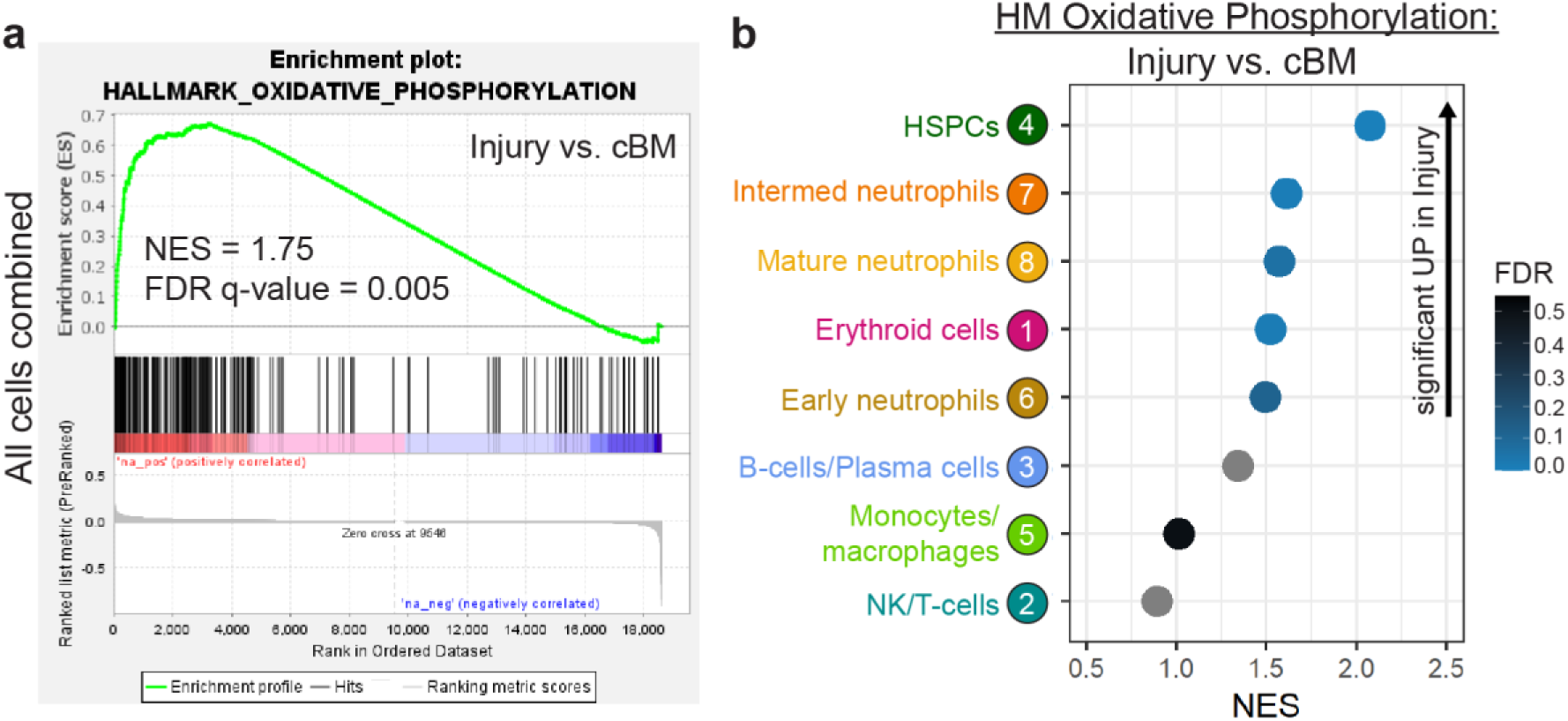
Analysis of genes relevant in oxidative phosphorylation. (**a**) GSEA on all cells combined (3 dpi): enrichment plot shows comparison with hallmark gene set oxidative phosphorylation indicating significant higher expression at the injury site when compared to the contralateral bone marrow. (**b**) GSEA separated for cell populations indicates that genes involved in oxidative phosphorylation are, particularly higher expressed in HSPCs, neutrophils, and erythroid cells.

**Extended Data Figure 8:**
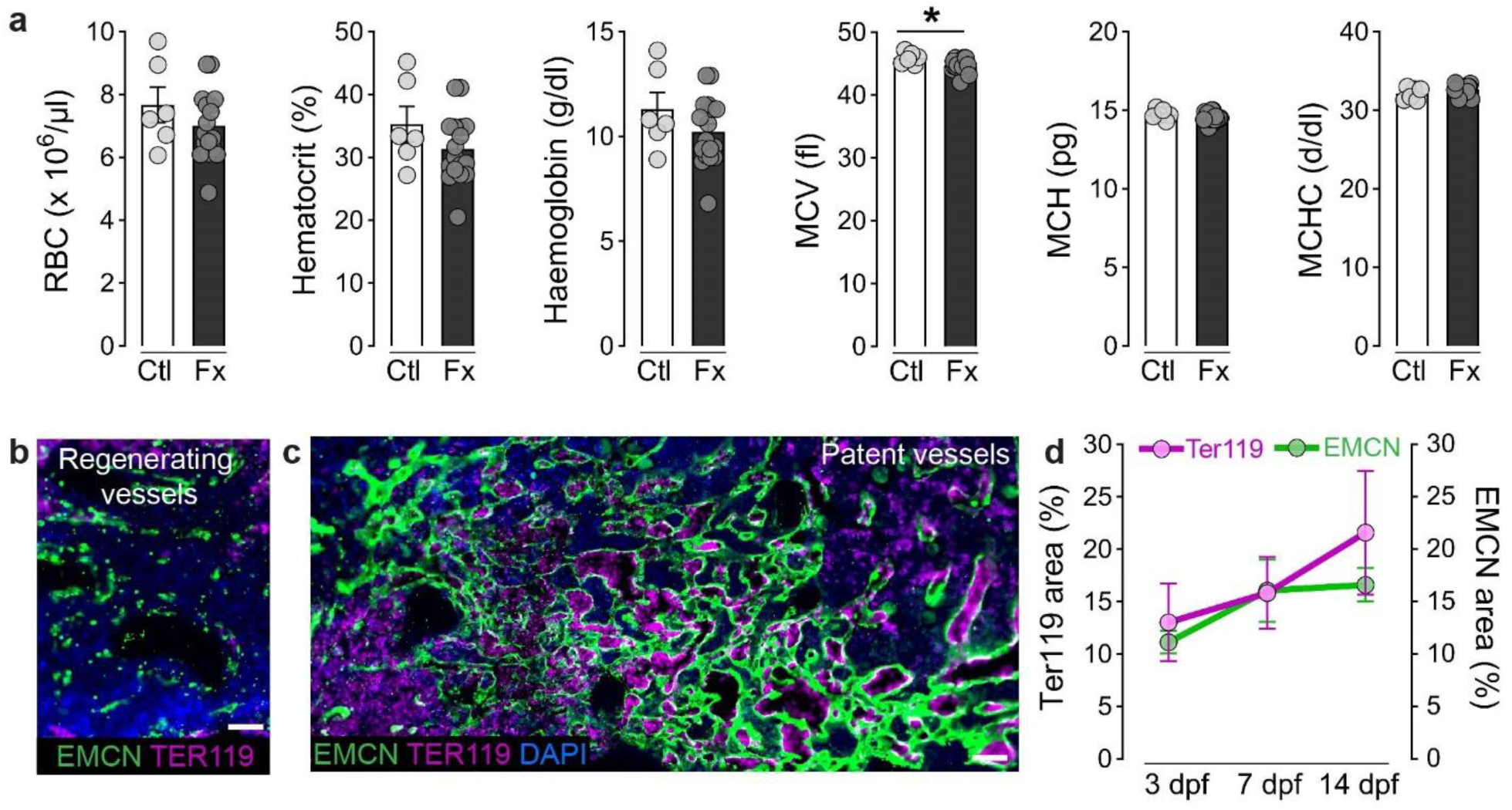
Bone injury activates local erythropoiesis at the injury site. (**a**) Blood parameters measured in peripheral blood at 3 dpi. Data show individual data points from n = 6-17 mice from more than 5 independent experiments and are mean ± s.e.m. *P* values were calculated using two-tailed Student’s *t*-test. \**P* < 0.05. (**b, c**) Representative images of Ter119 staining at the injury site at 3 dpi. (**b**) After injury, regenerating vessels do not contain Ter119^+^ erythrocytes compared to (**c**) patent vessels. Scale bars, 50 µm. Representative images for n = 4-8 mice per timepoint from 3 independent experiments. (**d**) Quantification of Ter119^+^ area and EMCN^+^ area at the injury site over time shows that the presence of erythrocytes depends on vascular regeneration. Data are mean ± s.e.m. for n = 4-8 mice per timepoint from 3 independent experiments. *P* values were calculated using one-way ANOVA with Tukey multiple comparison test.

**Extended Data Figure 9:**
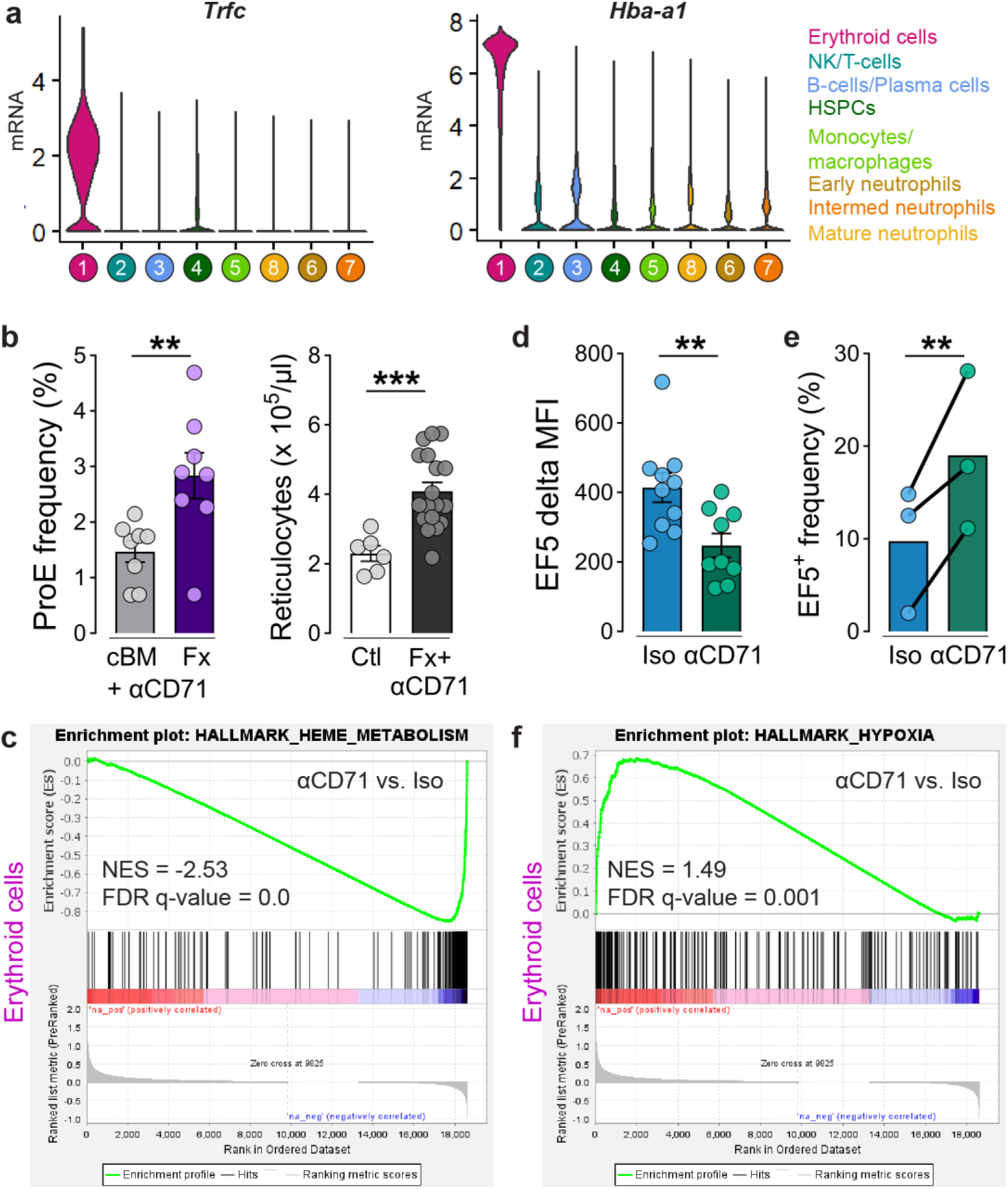
CD71 blockade does not affect erythropoiesis *per se* but induces hypoxia. (**a**) Expression of *Trfc* and *Hba-a1* across cell subpopulations highlights that erythroid cells express significantly higher levels of Transferrin receptor 1 (CD71, encoded by *Trfc*) than any other cell type, making them the primary target of CD71 antibody during injury in our experiments. Integrated data sets from n = 3 mice per group from 3 independent experiments (**b**) Frequency of ProE (Ter119^med^ CD71^high^) at injury site and concentration of reticulocytes in peripheral blood 3 dpi. Data show individual data points from n = 6-18 mice from more than 3 independent experiments and are mean ± s.e.m. *P* values were calculated using two-tailed Student’s *t*-test. (**c**) GSEA on erythroid cell cluster: enrichment plot shows comparison with hallmark gene set heme metabolism indicating significant lower expression with CD71 antibody treatment. (**d**) Quantification delta EF5 MFI to cBM including all cells. Data show individual data points for n = 8-9 mice per group from 5 independent experiments and are mean ± s.e.m. *P* values were calculated using two-tailed Student’s *t*-test. MFI, median fluorescence intensity. (**e**) Flow cytometry analysis of erythroid lineage cells (CD71^+^ Ter119^+^) and their EF5 staining intensity - representation of paired samples per experiment. Data show individual data points for n = 3-5 mice per group from 3 independent experiments and are mean ± s.e.m. *P* values were calculated using two-tailed paired Student’s *t*-test. (**f**) GSEA on erythroid cell cluster: enrichment plot shows comparison with hallmark gene set hypoxia indicating significant higher expression with CD71 antibody treatment. \*\**P* < 0.01 and \*\*\**P* < 0.001.

**Extended Data Figure 10:**
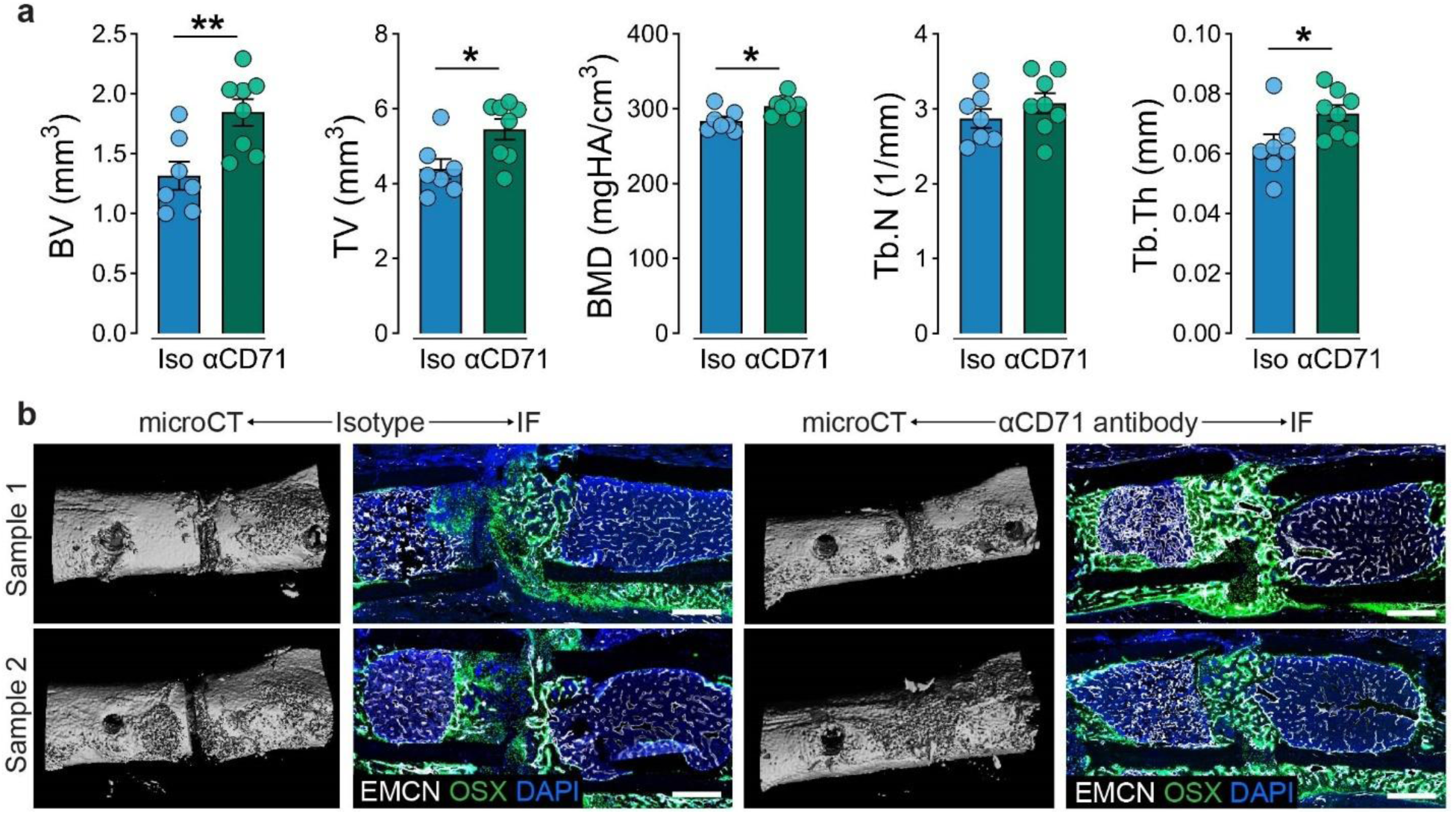
CD71 blockade promotes tissue regeneration at 14 dpi. (**a**) MicroCT analysis of bone morphometry. Data show individual data points for n = 9-10 mice per group from 2 independent experiments and are mean ± s.e.m. *P* values were calculated using two-tailed Student’s *t*-test. \**P* < 0.05, \*\**P* < 0.01 and \*\*\**P* < 0.001. (**b**) Additional samples: macroscopic 3D microCT reconstruction and corresponding immunofluorescence images of EMCN^+^ vessel formation and OSX^+^ progenitor infiltration at the injury site at 14 dpi. Scale bars, 500 µm.

**Extended Data Figure 11:**
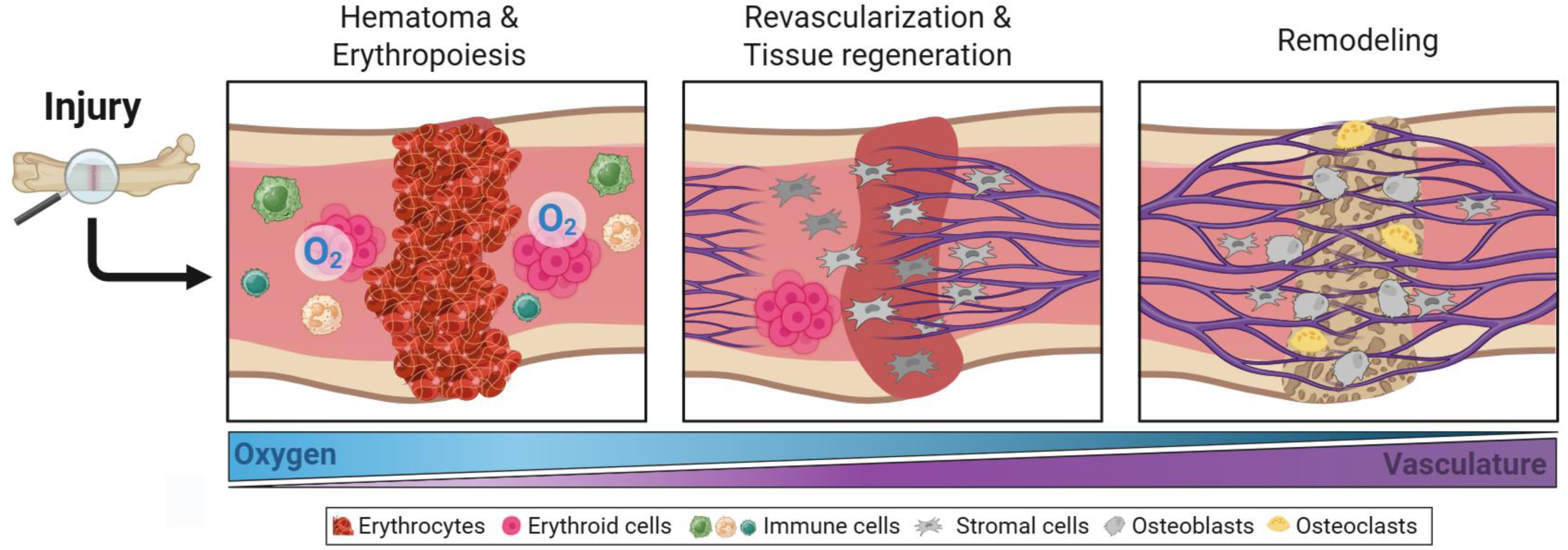
Illustration summarizing the role of erythroid cells during tissue regeneration in long bones. Upon injury/fracture, oxygen tension at the injury site increases due to bleeding into the hematoma and the onset of erythropoiesis in the adjacent bone marrow, during which erythroid precursor cells bind oxygen locally. Vascular regeneration proceeds inversely with oxygen tension, which gradually normalizes as healing progresses. Revascularization supports stem and stromal cell invasion, bone cell formation and activity which facilitates tissue regeneration, and ensures the reestablishment of the hypoxic bone marrow niche. Created in BioRender. Lang, A. (2025) https://BioRender.com/0r5qsqz.

